# Shotgun lipidomics and mass spectrometry imaging unveil diversity and dynamics in lipid composition in *Gammarus fossarum*

**DOI:** 10.1101/2020.12.01.368563

**Authors:** Tingting Fu, Oskar Knittelfelder, Olivier Geffard, Yohann Clément, Eric Testet, Nicolas Elie, David Touboul, Khedidja Abbaci, Andrej Shevchenko, Jerome Lemoine, Arnaud Chaumot, Arnaud Salvador, Davide Degli-Esposti, Sophie Ayciriex

**Affiliations:** Univ Lyon, CNRS, Université Claude Bernard Lyon 1, Institut des Sciences Analytiques, UMR 5280, 5 rue de la Doua, F-69100 Villeurbanne, France; Max Planck Institute of Molecular Cell Biology and Genetics, Pfotenhauerstraße 108, 01307 Dresden, Germany; INRAE, UR RiverLy, Ecotoxicology Team, F-69625 Villeurbanne, France; Laboratoire de Biogenèse Membranaire (LBM), CNRS, University of Bordeaux, UMR 5200, F-33882 Villenave d’Ornon, France; Université Paris-Saclay, CNRS, Institut de Chimie des Substances Naturelles, UPR 2301, 91198, Gifsur-Yvette, France

**Author notes:** Chemistry department, Humboldt University of Berlin, Brook-Taylor-Straße 2, 12489 Berlin (Oskar Knittelfelder).

**Keywords:** Shotgun lipidomics, lipidome, *Gammarus fossarum*, mass spectrometry imaging, sulfate-based lipids

## Abstract

Sentinel species are playing an indispensable role in monitoring environmental pollution in aquatic ecosystems. Many pollutants found in water prove to be endocrine disrupting chemicals that could cause disruptions in lipid homeostasis in aquatic species. A comprehensive profiling of the lipidome of these species is thus an essential step towards understanding the mechanism of toxicity induced by pollutants. We here extensively examined both the composition and spatial distribution of lipids in freshwater crustacean *Gammarus fossarum*. The baseline lipidome of gammarids of different gender and reproductive stage was established by high throughput shotgun lipidomics. Spatial lipid mapping by high resolution mass spectrometry imaging led to the discovery of sulfate-based lipids in hepatopancreas and their accumulation in mature oocytes. We uncovered in *G. fossarum* a diverse and dynamic lipid composition that deepens our understanding of the biochemical changes during development and which could serve as a reference for future ecotoxicological studies.

## INTRODUCTION

Lipids are a structurally diverse group of molecules that can be classified into several categories including fatty acyls, glycerolipids, glycerophospholipids, sphingolipids, sterol lipids, prenol lipids, saccharolipids, and polyketides according to the LIPID MAPS lipid classification system (Fahy, et al., 2011; Fahy, et al., 2005; Fahy, et al., 2009). Besides their basic function as building blocks of cell membrane, lipids are involved in essential biofunctions including signaling and energy storage that mediate cell growth, reproduction, and so on (Dutta and Sinha, 2017; Obeid, et al., 1993; van Meer, 2005; van Meer, et al., 2008). Lipid homeostasis in the organisms is extremely crucial for their development, maintenance and reproduction (De Mendoza and Pilon, 2019; Klose, et al., 2012; Zhang and Rock, 2008). As a result of anthropogenic activities, especially the intense use of chemical products, various kinds of pollutants are continuously released into the aquatic environment. Increasing evidence has shown that some of these pollutants are endocrine disrupting chemicals, also referred to as obesogens, which could interfere lipid homeostasis and cause toxic effects on a number of aquatic animal species (Capitao, et al., 2017; Fuertes, et al., 2020). Therefore, it is highly desirable to get retrospective and prospective comprehensive profiles of the lipidome in sentinel organisms in order to understand which and how the lipid species could be altered by chemical pollutants, and to further assess the ecological risk incurred by the contaminated aquatic environment.

Freshwater sentinel species *Gammarus fossarum* is one of the most represented amphipod crustaceans widespread across European inland aquatic habitats (Wattier, et al., 2020). Its broad distribution and sensitivity to a wide range of contaminants has made this keystone species a valuable model organism in ecotoxicology (Besse, et al., 2013; Chaumot, et al., 2015; Dedourge-Geffard, et al., 2009; Kunz, et al., 2010; Mehennaoui, et al., 2016; Wigh, et al., 2017). Endocrine effects (e.g., accelerated oocyte maturation, smaller vitellogenic oocytes, and decreased spermatozoon production) have also been observed in this species when exposed to endocrine disrupting chemicals in wastewater effluents (Schirling, et al., 2005) and in laboratory experiments (Trapp, et al., 2015). It is thus of great interest to characterize the lipidome of this organism to understand the molecular mechanism underlying these endocrine effects and to develop biomarkers for early stage risk assessment of obesogen contamination in freshwater systems. Initial assessment of lipid perturbation in *Gammarus fossarum* exposed to a growth regulator insecticide has been recently reported (Arambourou, et al., 2018). However, only a limited number of lipid classes and molecular species in this non model organism have been described (Arambourou, et al., 2018; Fu, et al., 2020; Kolanowski, et al., 2007). Our knowledge about the lipid composition especially the associated dynamics in this species is scarce.

Lipidomics *per se* covers a broad range of mass spectrometry (MS) workflows that aim to identify and quantify a great variety of lipid classes, including their molecular species in biological systems (Hsu, 2018; Hu, et al., 2019; Klose, et al., 2013; Shevchenko and Simons, 2010; van Meer, 2005; Wenk, 2005). In addition, when required, advanced lipid structural characterization (e.g., double bond and *sn*-positions) is also readily achievable *via* MS related developments such as ozonolysis (Brown, et al., 2011), UV-induced photodissociation (Bowman, et al., 2017; Brown, et al., 2011; Ryan, et al., 2017; Williams, et al., 2017) and ion mobility spectrometry (Groessl, et al., 2015; Jackson, et al., 2008; Kim, et al., 2009; Leaptrot, et al., 2019). The most popular analytical platforms for lipidomic analysis are electrospray ionization (ESI)-MS based shotgun lipidomics (i.e., direct infusion MS) and liquid chromatography (LC)-MS. In contrast to the necessity of time consuming chromatography separation in LC-MS, shotgun lipidomics is a high-throughput approach which relies on the direct infusion of a crude lipid extract into the ion source of a mass spectrometer (Han and Gross, 2003; Han and Gross, 2005; Han and Gross, 2005; Hsu, 2018). The maintenance of a constant concentration of the delivered lipid extract provides a stable ionization environment thus enabling reproducible qualitative and quantitative detection of hundreds of molecular lipid species in a single run. Nowadays, high-resolution mass spectrometry (e.g., Fourier transform ion cyclotron resonance and Orbitrap) is frequently used in shotgun lipidomics and has tremendously increased the confidence of analysis with its capability of resolving isobaric lipid species (Zullig and Kofeler, 2020). Up to now, shotgun lipidomics has been successfully applied to describe the lipidome of a variety of biological systems like yeast cells (Ejsing, et al., 2009; Klose, et al., 2012), Drosophila (Carvalho, et al., 2012; Palm, et al., 2012), flatworm (Thommen, et al., 2019), nematode *Caenorhabditis elegans* (Penkov, et al., 2010) and freshwater crustacean *Daphnia magna* (Taylor, et al., 2017).

Despite the access of the molecular complexity and the identification of hundreds to thousands of chemical species offered by shotgun lipidomics, the spatial distribution of the measured molecular lipid species is missing due to the mandatory lipid extraction procedures. Even though a global evaluation of lipid content in an organism proves to be very valuable for studying the lipid metabolism variations associated with development or disease (Ayciriex, et al., 2017; Carvalho, et al., 2012; Guan, et al., 2013; Knittelfelder, et al., 2018; Wang, et al., 2020), the spatial distribution of these biomolecules is crucial to understand their modes of action in particular functional compartments. In the last two decades, mass spectrometry imaging has emerged as a novel tool to localize various molecules such as metabolites, lipids, drugs and so on in biological tissues without the need of labelling (Davoli, et al., 2020; McDonnell and Heeren, 2007; Spengler, 2015). By using a laser or an ion beam to generate ions directly from the tissue, MSI preserves the spatial localization of the ions and enables multiplexed molecular mapping of important structures in tissue sections. Among the various MSI techniques, secondary ion mass spectrometry (SIMS) is well recognized for its high spatial resolving power providing a micron or even submicron routine resolution (Ayciriex, et al., 2011; Benabdellah, et al., 2010; Touboul and Brunelle, 2016). Therefore, SIMS remains popular in biological imaging despite the severe molecular fragmentation due to the employment of energetic primary ion beams. Matrix assisted laser desorption/ionization (MALDI), on the other hand, enables intact molecular detection with a good spatial resolution typically >10 μm (Benabdellah, et al., 2010; Gessel, et al., 2014). Both SIMS and MALDI MS imaging techniques have been intensively employed for lipid mapping (Bich, et al., 2014; Djambazova, et al., 2020; Sämfors and Fletcher, 2020; Zemski Berry, et al., 2011).

With the aim to provide an exhaustive characterization of the lipidome of freshwater sentinel species *G. fossarum* and to disclose the lipid dynamics during the development, we performed shotgun lipidomics and mass spectrometry imaging on gammarids of different gender and distinct female reproductive stages. The baseline lipidome of this organism was defined by high throughput shotgun lipidomics using a robotic chip-based nano-ESI infusion device coupled to a high-resolution mass spectrometer. To reveal the *in-situ* localization of a variety of lipids in the organs and tissues, gammarid tissue sections were examined globally by MALDI MSI and scrutinized in detail by time of flight (TOF)-SIMS imaging. Several unknown sulfate-based lipids were uncovered in this organism and localized in the epithelium of hepatopancreas by high resolution SIMS imaging which subsequently guided the targeted high mass resolution analysis of hepatopancreas lipid extract for molecular identification and structural characterization. Dynamic distribution of these sulfate-based lipids in the course of reproduction or oocyte maturation was then investigated by mapping the oocytes of female gammarids at two different reproductive stages. Overall, our results provide for the first time both compositional and spatial information of the lipids in this crustacean species.

## RESULTS AND DISCUSSION

### Lipid composition of the *Gammarus fossarum* lipidome

Shotgun Lipidomics analyses *via* high-resolution mass spectrometry in positive and negative ion mode were conducted to decipher the lipidome of males and females of the freshwater crustacean *G. fossarum* and at specific female reproductive stages (Figure S1). The reproduction cycle of female gammarids (oogenesis/vitellogenesis, embryogenesis) is closely synchronized with molting and is now well characterized (Chaumot, et al., 2020; Geffard, et al., 2010; Schirling, et al., 2004). In total, six molt stages are defined according to the phenotypic features of the females, namely postmolt (A, B), intermolt (C1, C2) and premolt (D1, D2). Contrary to females, spermatogenesis in male gammarids is not related to the molting cycle and morphological parameters are not available to depict accurately the organisms at different spermatogenesis stages. In our study, all the male organisms sampled were in amplexus to ensure they were at similar spermatogenesis stage (mature). Female gammarids were collected at the beginning of intermolt stage (C1) and at premolt stage (D1) to investigate lipid alterations related to oocyte maturation.

By shotgun lipidomics, more than 200 molecular lipid species were quantified, corresponding to 11 major lipid classes in wild male and female adult gammarids – triacylglycerols (TAG), diacylglycerols (DAG), phosphatidylcholines (PC), phosphatidylethanolamines (PE), ether lipids (PE-O and PC-O), phosphatidylinositols (PI), lysophosphatidylcholines (LPC), lysophosphatidylethanolamines (LPE), sphingomyelins (SM), and cholesterol (Chol) (Figure 1A). Our findings have significantly expanded the lipidome coverage reported previously in *G. fossarum* in terms of both lipid class and molecular lipid species (Arambourou, et al., 2018). Total fatty acids (FA) including the acyl chains of larger lipid molecules from the whole organism were also examined and deciphered by GC-FID (Figure S2). The predominant fatty acid in gammarids was monounsaturated C18:1 (~33%). Other fatty acids of comparatively high level were saturated C16:0 (~16%), polyunsaturated C20:5 n-3 (~14%) and monounsaturated C16:1 (~13%). Omega-3 fatty acids made up ~25% of the polyunsaturated fatty acid (PUFA) in contrast of omega-6 PUFA (~9%). The main n-3 FA was eicosapentaenoic acid (20:5, n-3), followed by α-linolenic acid (FA 18:3, n-3).

**Figure 1.**
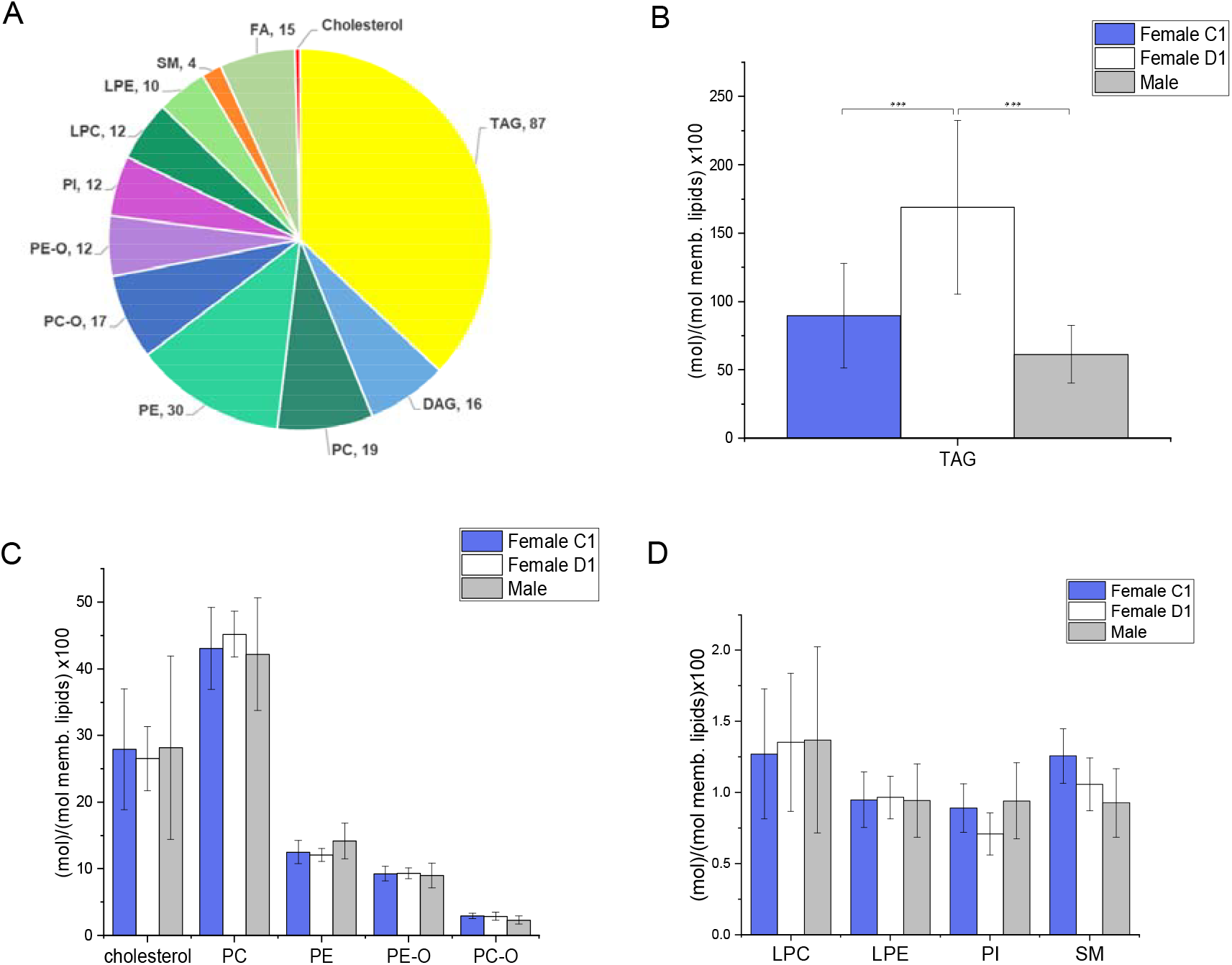
Lipid composition of the *Gammarus fossarum* lipidome. (A) Lipid classes identified in *G. fossarum* organism with the associated number of lipid species identified. Comparison of the different lipid profile between male and female gammarids at specific female reproductive stages (C1 versus D1): (B) TAG profile. (C) cholesterol, phosphatidylcholine, phosphatidylethanolamine and ether lipids distribution. (D) lysophospholipids, phosphatidylinositol and sphingomyelin profile. Lipid class abundance is presented as moles per mole of total membrane lipid (phospholipids, sphingolipids and sterols – not including storage lipids). Statistically significant changes (p*<*0.05) are marked by asterisks. Error bars indicate standard deviation.

TAG class turned out to be the most predominant lipid class in gammarids. Interestingly, female at D1 stage contained more TAG than female at C1 stage and male (Figure 1B), whereas no major changes were observed for membrane lipids like cholesterol, phospholipids (PC, PE, PE-O, PC-O, PI), lysophospholipids (LPE, LPC) and sphingomyelin (Figure 1C, 1D). In the reproductive cycle of female gammarids, D1 stage follows the second vitellogenesis where dramatically increased follicular surface has been observed compared to females at C1 postmolt stages (Geffard, et al., 2010). It is well documented that in many crustacean species, ovarian maturation requires huge amount of lipids to realize the vitellogenesis process (Alava, et al., 2007; Lee and Walker, 1995; Ravid, et al., 1999). In the female gammarids at D1 stage, TAG probably serves as energy storage reservoir (FA store) that might be rapidly mobilized on demand during the second vitellogenesis (Subramoniam, 2011). Although a global lipid accumulation during oocyte maturation is evident, the mostly affected stages and lipid species seem to differ among the crustacean species. Equal amounts of TAG and phospholipids are accumulated in the ovary of *Penaeus semisulcatus* when oocytes reach maturation (Ravid, et al., 1999). However, in most species, TAG is primarily responsible for the changes in total lipid content and increase of phospholipids only occurs at the end of maturation (Mourente, et al., 1994; Wouters, et al., 2001) as probably in the case of *G. fossarum*.

### Diversity of molecular lipid species in *G. fossarum* across gender and distinct female reproductive stages

Next, we questioned whether the profile of the molecular lipid species of each lipid class found in adult gammarid varies between the gender and different female reproductive stages. Figure 2 displays the profiles of the molecular species of four lipid classes (namely TAG, PC, PE and SM) in male gammarid and female gammarids at C1 and D1 stages. The abundance of each lipid was normalized to the total abundance of the corresponding lipid class to show the proportion of each molecular lipid species within a lipid class. It is observed that TAG species containing relatively short chain and poly-unsaturated fatty acids (e.g., TAG 46:1, TAG 46:2 and TAG 48:1 to TAG 48:6) have a higher proportion in male compared to female, whereas TAGs with longer chain and poly-unsaturated fatty acids are more prominent in female (e.g., TAG 54:6, TAG 56:7). This difference in the proportion of each molecular species in total TAG is less significant between the females at C1 and D1 reproductive stages. All the reported 46 TAG molecular species were characterized by tandem mass spectrometry. TAG precursors were detected in positive ion mode as ammonium adduct [M+NH_4_]^+^, of which the MS/MS spectra were featured by neutral losses (NLs) of NH_3_ and an acyl side chain (as a carbolic acid ROOH) to generate a diacyl product ion (Figure S3). For instance, under HCD, the precursor ion at *m/z* 868.7416 (TAG 52:6, [M+NH_4_]^+^), exhibited NLs of 319, 271, 299 and 245 which correspond to FA 20:5, 16:1, 18:1 and 14:0, respectively. We were able to determine that TAG 52:6 was composed of two isomers TAG 16:0-16:1-20:5 and 14:0-18:1-20:5 (Figure S3B).

**Figure 2.**
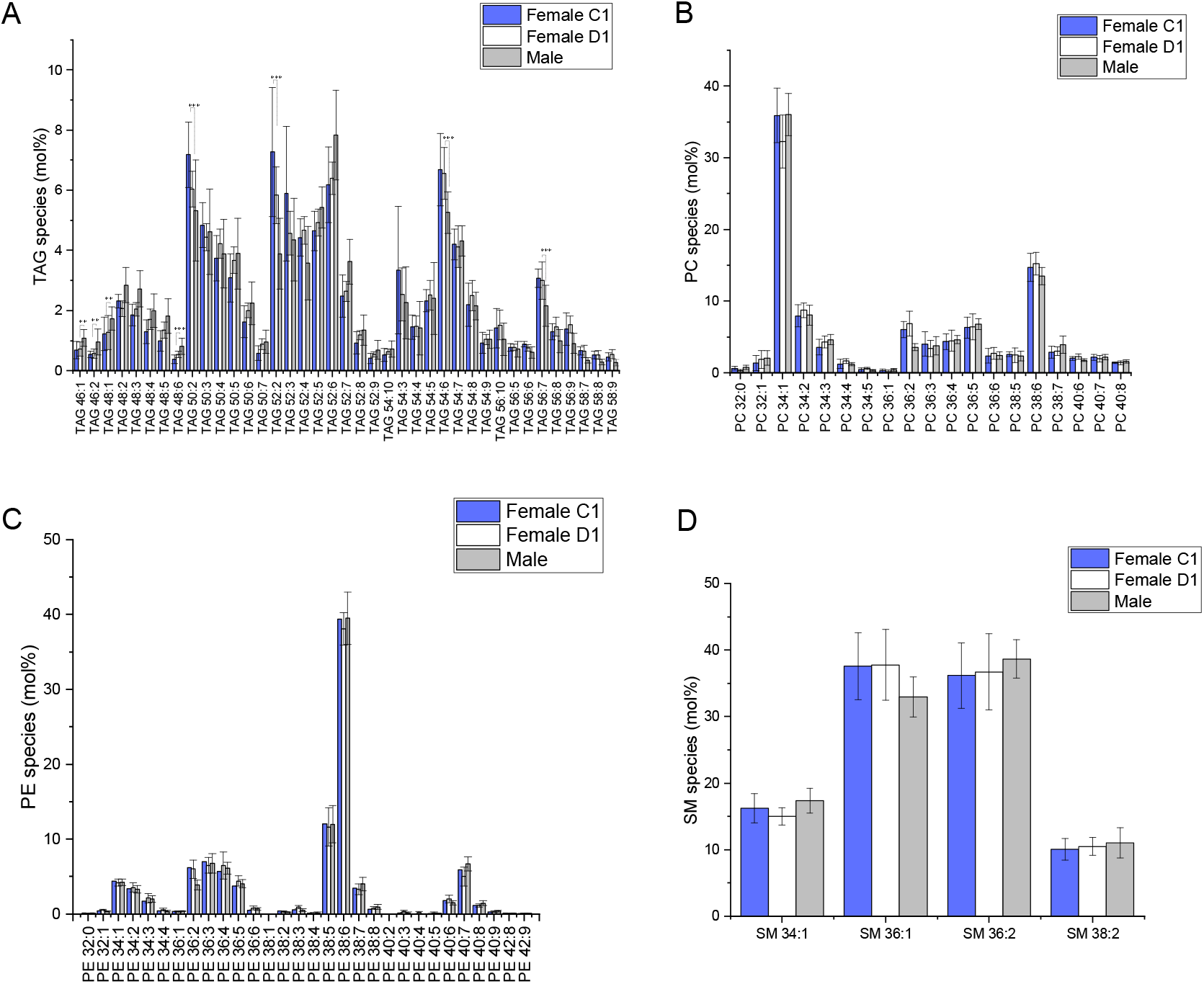
Diverse molecular lipid species in *G. fossarum* across gender and female reproductive stages. Lipid profile for the main glycerolipid (A) TAG and the two main phospholipid classes (B) PC and (C) PE. (D) Sphingomyelin profile. Note that for TAG profile only the species higher than 5 mol% are presented. Statistically significant changes (p<0.05) are marked by asterisks. Error bars indicate standard deviation.

For glycerophospholipid class, the predominant PC species in gammarids is monounsaturated PC 34:1, followed by the poly-unsaturated PC 38:4 (Figure 2B). These two major PC species were defined as PC 16:0-18:1 and PC 18:1-20:5, respectively, based on the observation of the fragments corresponding to fatty acids in the negative MS/MS spectra (Figure S4). For PE lipid class, PE 38:6 (PE 18:1-20:5) stand out as the most abundant molecular lipid species (Figure 2C and Figure S5). Only 4 SM species were detected in the gammarid, namely SM 34:1 (d18:1/16:0), 36:1 (d18:1/18:0), 36:2 (d18:1/18:1) and 38:2 (d18:1/20:1) (Figure 2D and Figure S6). This finding was confirmed by a targeted lipidomics approach employing LC-ESI-MS/MS (data not shown). Overall, no significant differences in the proportional abundance of the glycerophosholipid and sphingolipid lipids were observed between male and female gammarid at the molecular species level.

### Localization of lipids in whole-body gammarid section by MALDI MSI

With the rich molecular lipid species information obtained with shotgun lipidomics, we then performed mass spectrometry imaging to map the lipids *in situ* and reveal their spatial localization in the tissues and organs of *G. fossarum*. First, longitudinal sections of the male gammarid were mapped by MALDI MSI to examine the global distribution of the lipid species. In negative ion mode, the ions at *m/z* 295.2 and 297.2 were the main species detected from the tissue section (Figure S7), while in positive ion mode phosphatidylcholine (PC) were the prominent lipid species observed in the mass range of *m/z* 750-850 (Figure S8). MALDI MS ion images of the main PC lipids, PC 34:1 and PC 36:3, are displayed in Figure 3. Both PC species are distributed across the whole body in cephalon, muscle and thorax segments (TS) with higher abundance in cephalon. PC 34:1 is the most abundant PC species according to the shotgun analysis (Figure 2). Three ions related to PC 34:1 ([M+H]^+^, [M+Na]^+^ and [M+K]^+^) were detected, all showing identical distribution in the cephalon, muscle and TS tissue of the male gammarid. Besides the PC lipids, an ion at *m/z* 841.5 was also observed in the measured mass range (Figure 3F and Figure S8). The ion image showed a distinct spatial localization from the PC lipids. By overlaying the ion image with that of PC 34:1 [M+K]^+^ and subsequently the optical image of the analyzed tissue section (Figure 3G and 3I), it is revealed that this molecule is principally colocalized to the gonad as well as to the area close to hepatopancreas (HP) where the gonad is usually located but not seen in the optical image due to the non-ideal cutting orientation during cryosectioning. This ion at *m/z* 841.5 was tentatively attributed as sulfated glyceroglycolipid (SGG), also referred to as seminolipid with C16:0/C16:0 alkyl/acyl chains (Lessig, et al., 2004).

**Figure 3.**
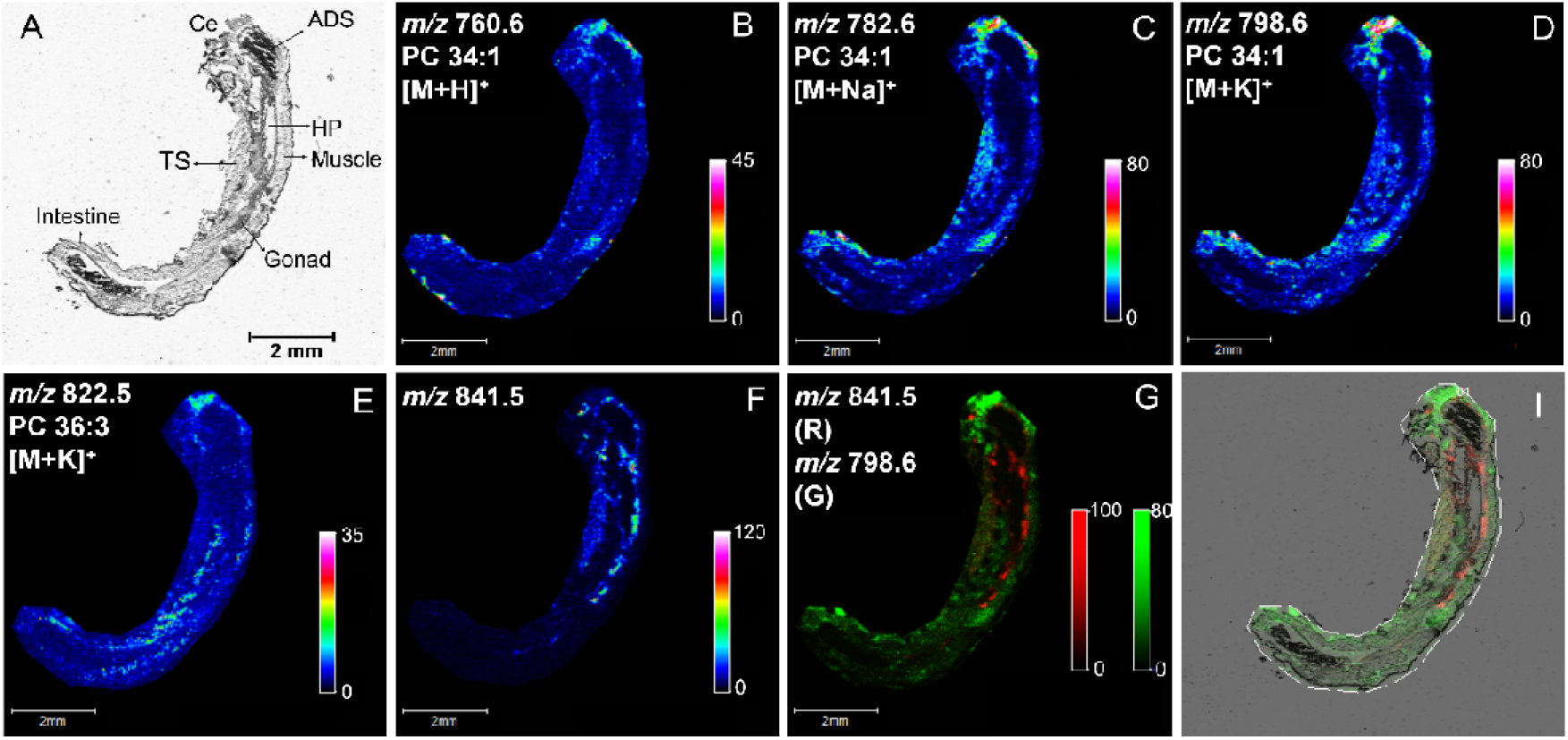
Localization of lipids in whole-body gammarid section by MALDI MSI. (A) Optical image of the whole-body tissue section of the male gammarid. (B) Ion image of protonated PC 34:1. (C) Ion image of sodium adduct of PC 34:1. (D) Ion image of potassium adduct of PC 34:1. (E) Ion image of potassium adduct of PC 36:3. (F) Ion image of the ion at *m/z* 841.5. (G) Two-color overlay of the ion at *m/z* 841.5 and the potassium adduct of PC 34:1. (I) Overlay of the two-color overlay image in G and the optical image. Ce: Cephalon; ADS: anterior digestive system; HP: hepatopancreas; TS: thorax segments.

Seminolipid C16:0/16:0 is the predominant SGG species and is a key lipid involved in germ cell differentiation during spermatogenesis in mammalians (Tanphaichitr, et al., 2018; Zhang, et al., 2005), and it is likely to have similar function in *G. fossarum*. In the anatomy of male gammarid, the gonad is surrounded by orange lipid droplets (Wigh, et al., 2017). Therefore, it is unclear if this molecule detected here is derived from the gonad tissue or from the surrounding lipid droplets. Our future work will be focusing on characterizing this seminolipid, discovering its potential analogues and identifying its precise localization.

### Lipid distribution in targeted organs by high resolution SIMS imaging

After examining the global distribution of lipids in the male gammarid, we then applied high resolution SIMS imaging to scrutinize the individual organs at 2 μm lateral resolution. Two regions of interest (ROIs) covering various tissue types including hepatopancreas, gonad and muscle were targeted as shown in the optical images in Figure 4. The ion at *m/z* 224.1 which is a characteristic fragment of phosphatidylcholine (PC) lipids was found abundant in the muscle tissue (Figure 4B and 4D), consistent with the results from MALDI MSI. It is interesting to note that this PC fragment was also observed with high intensity in the gonad tissue, implying the presence of PC lipids in the gonad and which may differ from those in the muscle as certain PC species such as PC 34:1 was not observed in the gonad from MALDI MSI analysis. Vitamin E was also observed in SIMS MSI analysis and was found in all kinds of tissue types. Several unknown ion species were detected between *m/z* 648.4 and *m/z* 696.4 in the positive ion mode (Figure S9) and the corresponding ion images show that they appear to be specifically colocalized to the hepatopancreas (Figure 4B and 4D).

**Figure 4.**
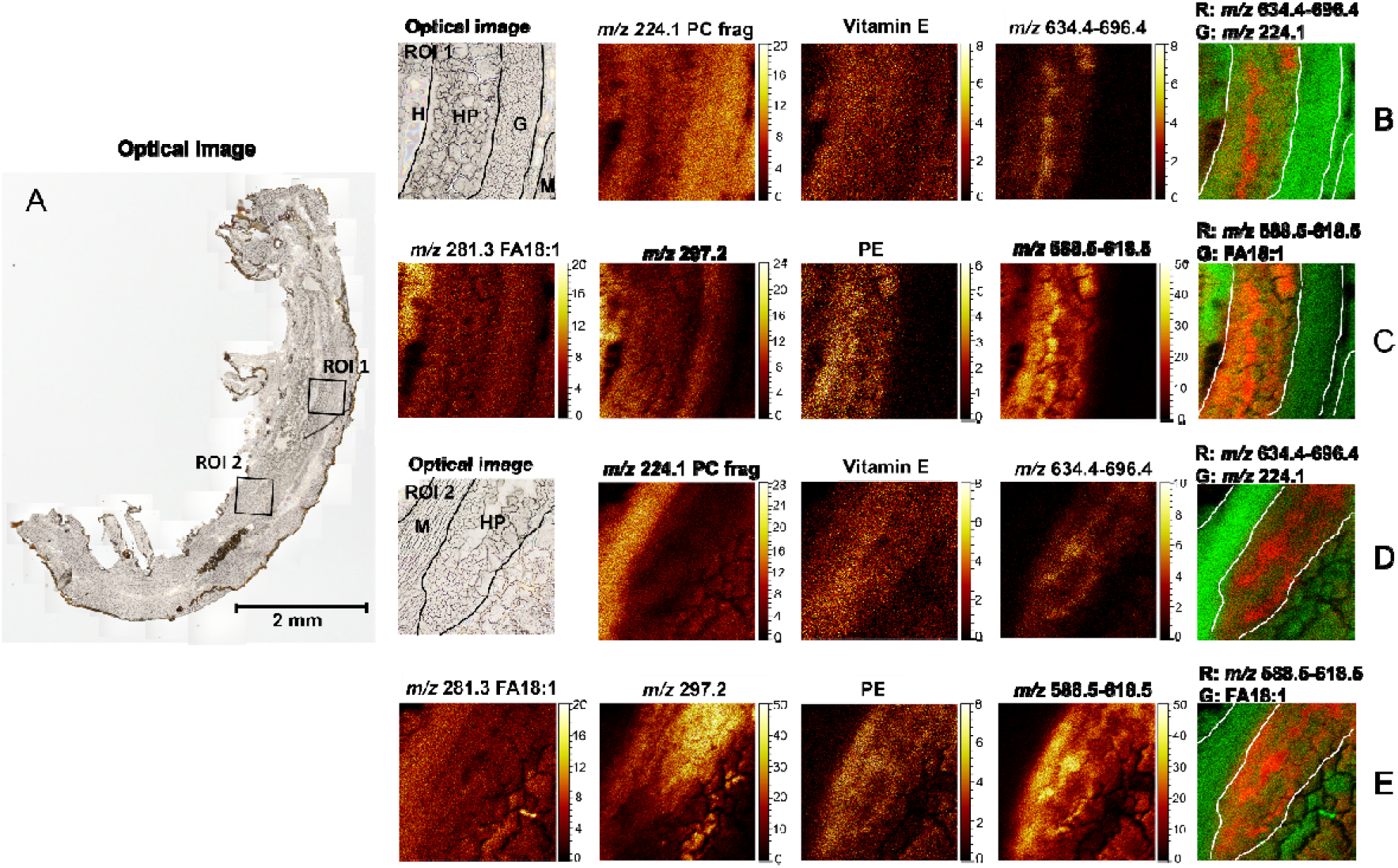
Lipid distribution in targeted organs of male gammarid by high resolution SIMS imaging. ROI: region of interest. (A) Optical image of the gammarid tissue section. (B) Ion images of lipid species detected in positive ion mode in ROI 1. (C) Ion images of lipid species detected in negative ion mode in ROI 1. (D) Ion images of lipid species detected in positive ion mode in ROI 2. (E) Ion images of lipid species detected in negative ion mode in ROI 2. H: haemocoel. HP: hepatopancreas. G. gonad. M: muscle. Ion image of Vitamin E was summed from those of ions at *m/z* 429.4 and *m/z* 430.4. Ion image of *m/z* 634.4-696.4 was summed from those of ions at *m/z* 634.4, 648.5, 648.4, 664.5, 666.4, 680.4, 682.4 and 696.4. Ion image of *m/z* 588.5-618.5 was summed from those of ions at *m/z* 588.5, 602.5, 604.5, 616.5 and 618.5.

In negative ion mode, several fatty acid species were detected (Figure S10), among which FA 18:1 (oleic acid) was observed across the analyzed regions with higher abundance observed in the body cavity haemocoel (Figure 4C and 4E). The ion at *m/z* 297.2 showed strong signal in the SIMS spectra acquired in negative ion mode and the ion images illustrate that this ion species seems to be mostly derived from the lumen of the hepatopancreas (Figure 4E). The ion image of phosphatidylethanolamine (PE) lipid was summed from those of diacylglycerophosphatidylethanolamine and 1-(1Z-alkenyl),2-acylglycerophosphatidylethanol-amines which were all predominantly localized in the hepatopancreas (Figure S11). Also in hepatopancreas some unknown ion species at *m/z* 588.5, 602.5, 604.5, 616.5 and 618.5 were detected. These ions turned out to have a very different distribution pattern from that of the ion at *m/z* 297.2, although they were all derived from the hepatopancreas organ (Figure S12, Figure 4C and 4E). The overlay images of the sum of these ion species and FA 18:1 reveal that their distribution is similar to that of ions at *m/z* 648.4-696.4 detected in positive ion mode. This high-resolution examination of regions of interest of the whole-body tissue section provided not only a clearer map of the chemical species distributed in individual organs of the gammarid, but also supplementary information in terms of lipid detection.

### Identification of sulfate-based lipid species in hepatopancreas

With the aim to identify whether the ion species at *m/z* 632.4-696.4 (in positive ion mode) and *m/z* 588.5-618.5 (in negative ion mode) are localized in the lumen or the epithelium of the hepatopancreas, we then targeted the hepatopancreas region by SIMS imaging on the transverse section where the structures of the 4 hepatopancreas caeca are well defined (Figure 5). Consistent with the above MALDI and SIMS analyses of longitudinal tissue sections, the ion at *m/z* 224.1 (PC head group) was predominantly found in the muscle tissue. The ion at *m/z* 196.9 which was assigned as salt ion K_2_NaSO_4_^+^ (based on spectral library search in SurfaceLab) turned out to be concentrated in the lumen of the hepatopancreas, whereas the ions at *m/z* 632.4-696.4 were mainly detected from the epithelium of the hepatopancreas and intestine (Figure 5A). For ions generated in negative ion mode, FA 18:1 shared similar distribution as PC head group. The ion at *m/z* 297.2 was found in both the lumen and epithelium of hepatopancreas with higher abundance in the lumen. The summed ion image of the ions at *m/z* 588.5-618.5 and its overlay with FA 18:1 and *m/z* 297.2 confirm that these ion species are localized principally in the epithelium of hepatopancreas and intestine (Figure 5B).

**Figure 5.**
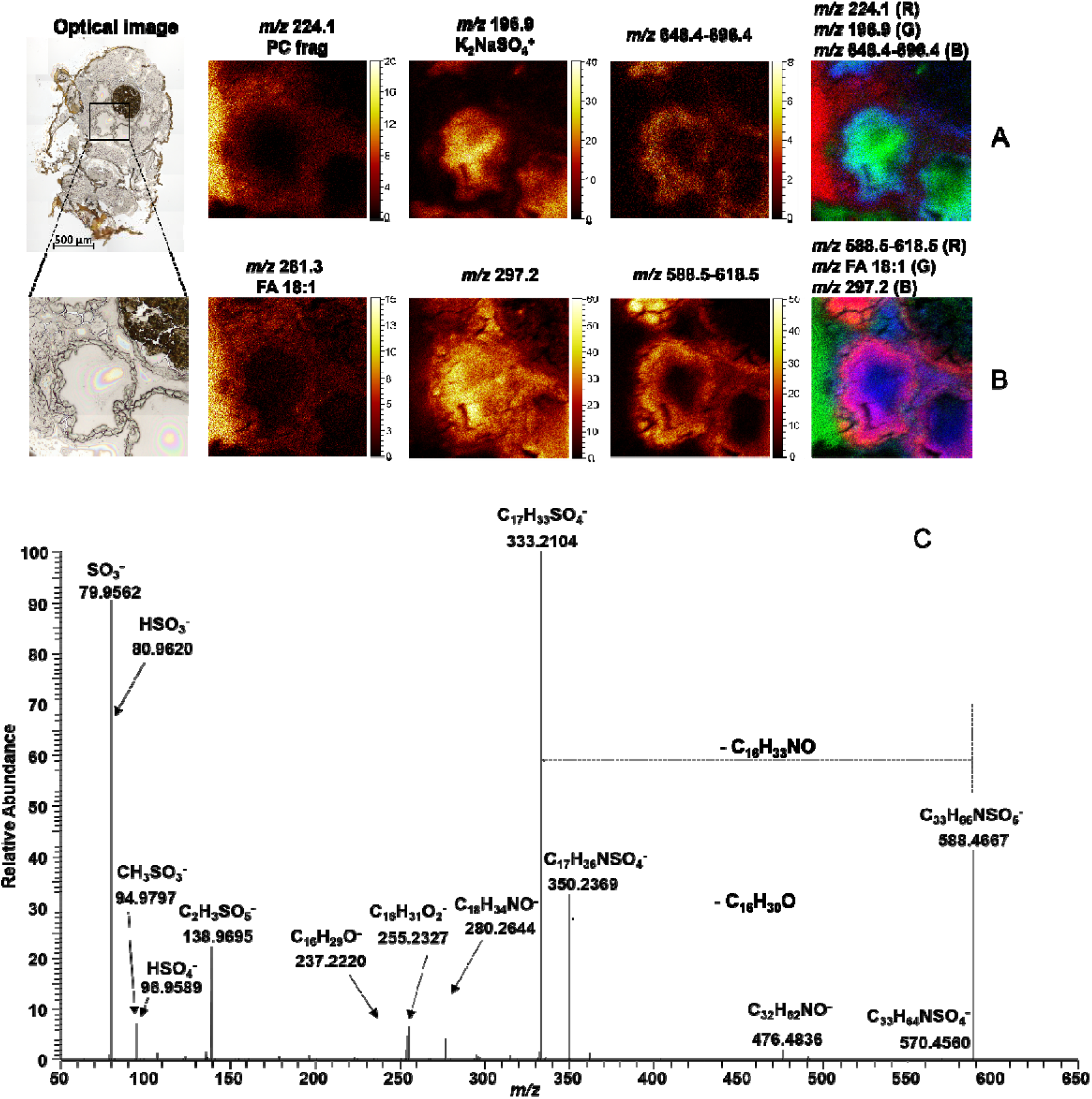
Identification of sulfate-based lipid species in hepatopancreas. (A) Ion images of selected chemical species detected in positive ion mode. (B) Ion images of selected chemical species detected in negative ion mode. (C) MS/MS spectrum of the precursor ion at *m/z* 588.4667 acquired on a Q Exactive mass spectrometer in negative ion mode.

To characterize these unknown ion species, hepatopancreas tissues from 10 male gammarids were pooled and then extracted according to a modified Folch lipid extraction procedure. Full scan mode and MS/MS on a high resolution mass spectrometer (ESI-Orbitrap) was performed in both polarities. The ions at *m/z* 634.4-696.4 were not observed in the positive ion mode, either due to low abundance or low extraction efficiency. In negative ion mode, the ions at *m/z* 295.2, 297.2 and *m/z* 588.5-618.5 were readily detected and subsequent MS/MS analyses yielded informative tandem mass spectra (Figure 5C and Figures S13-S18). Fragment ions corresponding to sulfate ion (*m/z* 79.9562, SO_3_^−^, *m/z* 80.9620, HSO_3_^−^ and *m/z* 96.9589, HSO_4_^−^) as well as neutral losses of hydrocarbon chains containing O and N motifs (e.g., C_16_H_30_O and C_16_H_33_NO in Figure 5C) were observed in the MS/MS spectra of all the precursor ions at *m/z* 588.5-618.5, indicating these molecules were sulfate-based lipids. Both precursor and fragment ions were annotated with a high mass accuracy better than 5 ppm (Figure 5C, Figures S15-S18, and table S1). Based on these accurate MS and MS/MS data, we interrogated various databases including METLIN (https://metlin.scripps.edu/landing_page.php?pgcontent=mainPage) and LIPID MAPS (https://www.lipidmaps.org/) without getting any possible matches. Thus, these sulfate-based lipids are probably new molecules present in this gammarid species. Sulfate-based lipids (SL) are a subclass of sulfolipids which is a heterogenous group of lipids containing sulfur element (Dias, et al., 2019). In mammals, SL are involved in various biochemical processes including cell-cell communication (Honke, 2013), inflammation (Hu, et al., 2007) and immunity (Avila, et al., 1996). Some SL such as cholesterol sulfate and SO_3_-Gal-ceramide are commonly found in the epithelium of digestive tracts to regulate the activities of pancreatic protease by inhibiting elastase (Ito, et al., 1998). We hypothesized that the SL detected in the hepatopancreas might be involved in similar activities in *G. fossarum*. However, further investigations are required to elucidate their biological functions.

### Dynamic change in oocyte lipid composition during the female reproductive cycle

As described previously, the reproductive cycle of female gammarids comprises six molt stages which are characterized by the maturation of oocytes (Geffard, et al., 2010; Schirling, et al., 2004). To investigate the dynamics of lipid composition related to the maturation process, high resolution TOF-SIMS imaging was performed to map the chemical composition of the early vitellogenic oocytes (EVO) of female gammarids at C1 stage and the late vitellogenic oocytes (LVO) of female gammarids at D1 stage, respectively. The analyzed areas covering various tissue types including oocytes (Ooc), hepatopancreas (HP), intestine (In) and muscle are shown in Figure 6. The regions of oocytes were defined and outlined by comparing with H&E stained images of the same sections analyzed by TOF-SIMS (Figure S19). For females at both C1 and D1 stages, fatty acids were detected across the analyzed area with lower abundance in the intestine and hepatopancreas. Intense signals of fatty acids were also observed in the hallow area caused by tissue cracking which frequently occurred during preparation of fragile tissue sections. The ion at *m/z* 297.2 was found abundant in hepatopancreas and intestine in females at both stages. Very interesting to note is the distribution of the newly identified sulfate-based lipids. Compared to the female at C1 stage where the SL were mainly detected from the hepatopancreas, the D1 female showed a significant accumulation of SL in the oocytes. In Figure 6B, the overlay image of SL, FA18:1 and the ion at *m/z* 297.2 illustrates that the SL are only accumulated in the secondary oocytes (SO) and are absent in the primary oocytes (PO) which are immature oocytes at previtellogenic stage (Schirling, et al., 2004; Tan-Fermin and Pudadera, 1989).

**Figure 6.**
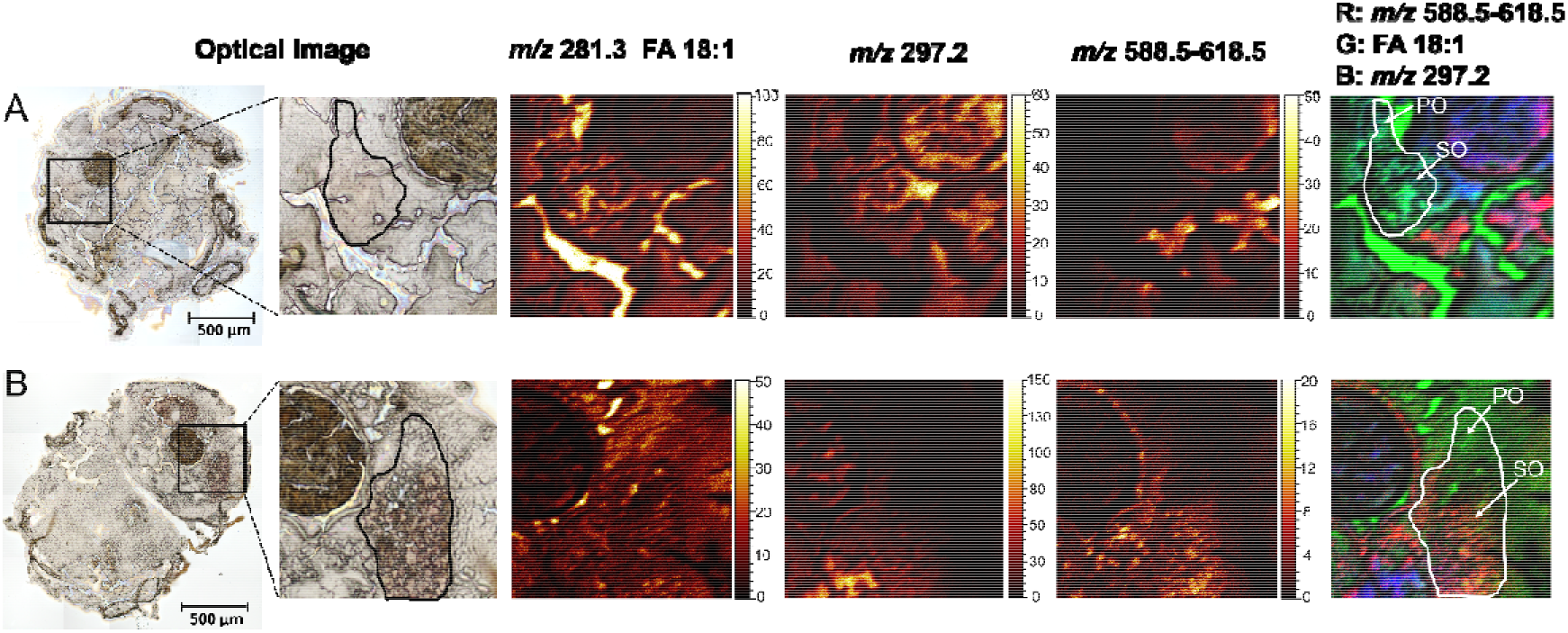
Dynamic change in oocyte lipid composition during the reproductive cycle. (A) Optical image of the transvers tissue section of C1 female and ion images of FA18:1 and sulfate-based lipids. (B) Optical image of the transvers tissue section of D1 female and ion images of FA 18:1 and sulfate-based lipids. The ion image of sulfate-based lipids at *m/z* 588.5-618.5 was summed from those of *m/z* 588.5, *m/z* 602.5, *m/z* 604.5, *m/z* 616.5 and *m/z* 618.5.

Although it is demonstrated in many crustacean species that accumulation of lipids in the oocytes occurs during the ovarian maturation, the origin of these rapidly accumulated lipids is not fully understood. Mobilization of lipids from hepatopancreas to oocytes in the prawn *Penaeus japonicus* has been proved by tracing the lipids derived from radioactive labelled fatty acids (Teshima, et al., 1988). In addition, several studies have reported the co-occurrence of a decreased lipid content in hepatopancreas and an increased lipid content in the ovary during ovarian maturation (Alava, et al., 2007; Castille and Lawrence, 1989; Spaargaren and Haefner, 1994). Therefore, it has been hypothesized that hepatopancreas also functions as lipid storage organ in crustacean species and could release the required lipids to oocytes to facilitate their maturation. By high resolution mass spectrometry imaging, our observation of the accumulation of SL in oocytes of D1 female suggests that these SL have probably gone through this lipid transfer process to accumulate in oocytes of the female gammarids and play an important role in the maturation process.

### Significance

The comprehensive lipid profile of *G. fossarum* revealed by shotgun lipidomics and mass spectrometry imaging enables us to obtain in-depth insights into the lipid homeostasis in this sentinel crustacean species. Guided by high resolution mass spectrometry imaging, the discovery of sulfate-based lipids in the epithelium of the hepatopancreas and intestine indicates a molecular similarity and very probably a functional similarity as well between the digestive tracts of this species and mammals. We also showcased a dynamic lipid composition in the oocytes during the reproductive cycle, which supports the hypothesis that lipid transfer from hepatopancreas to oocytes could occur to provide necessary lipids for the oocyte maturation. In addition, this exhaustive lipid profile of sentinel species *G. fossarum* will serve as a valuable reference for future investigations in the disruptions of lipid homeostasis caused by environmental stressor exposure. Thus, the application of the presented methodology in environmental relevant species could contribute to providing molecular mechanisms of the observable toxic effects of the contaminants and to improving our understanding of ecosystem health alterations.

## STAR METHODS

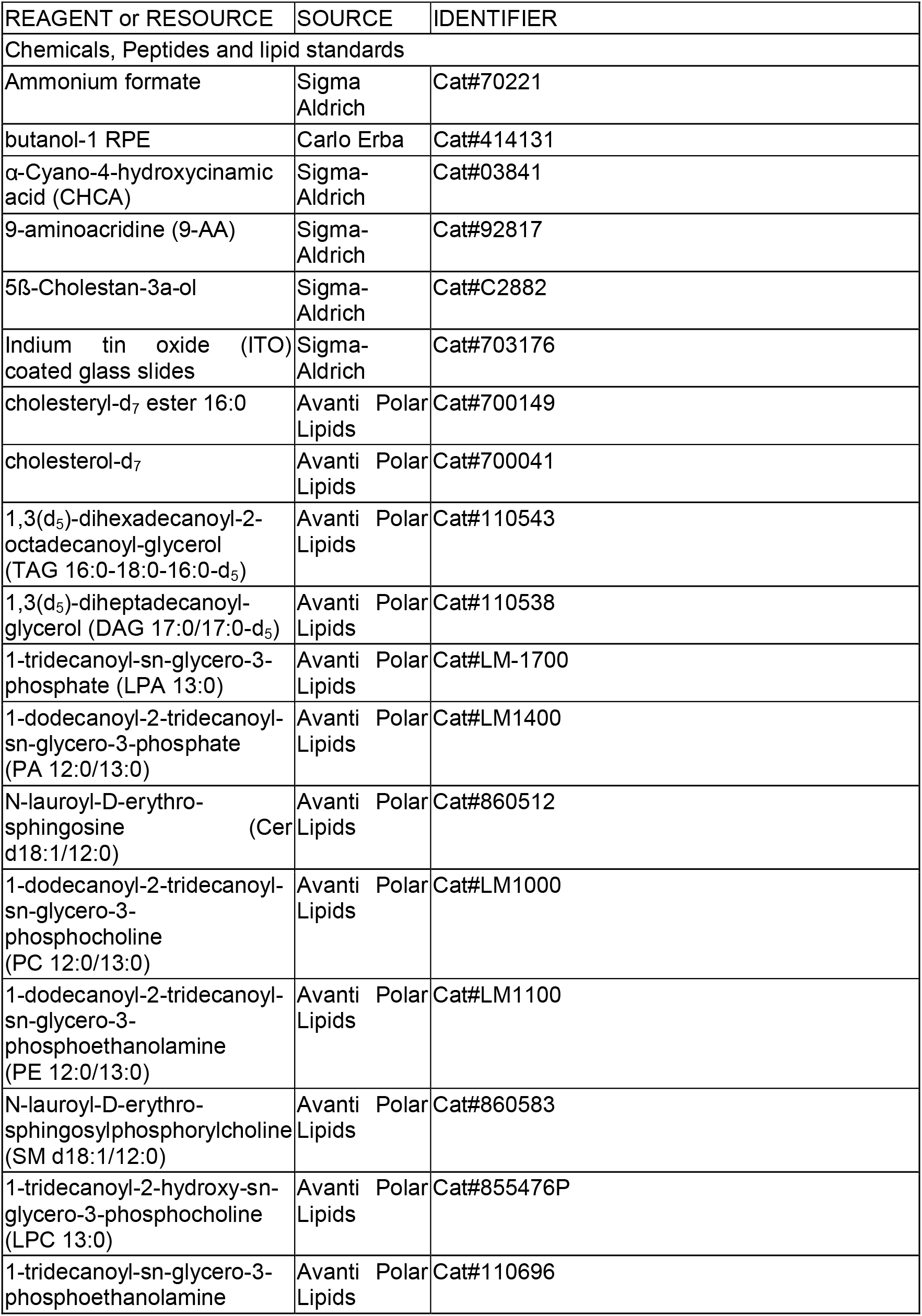
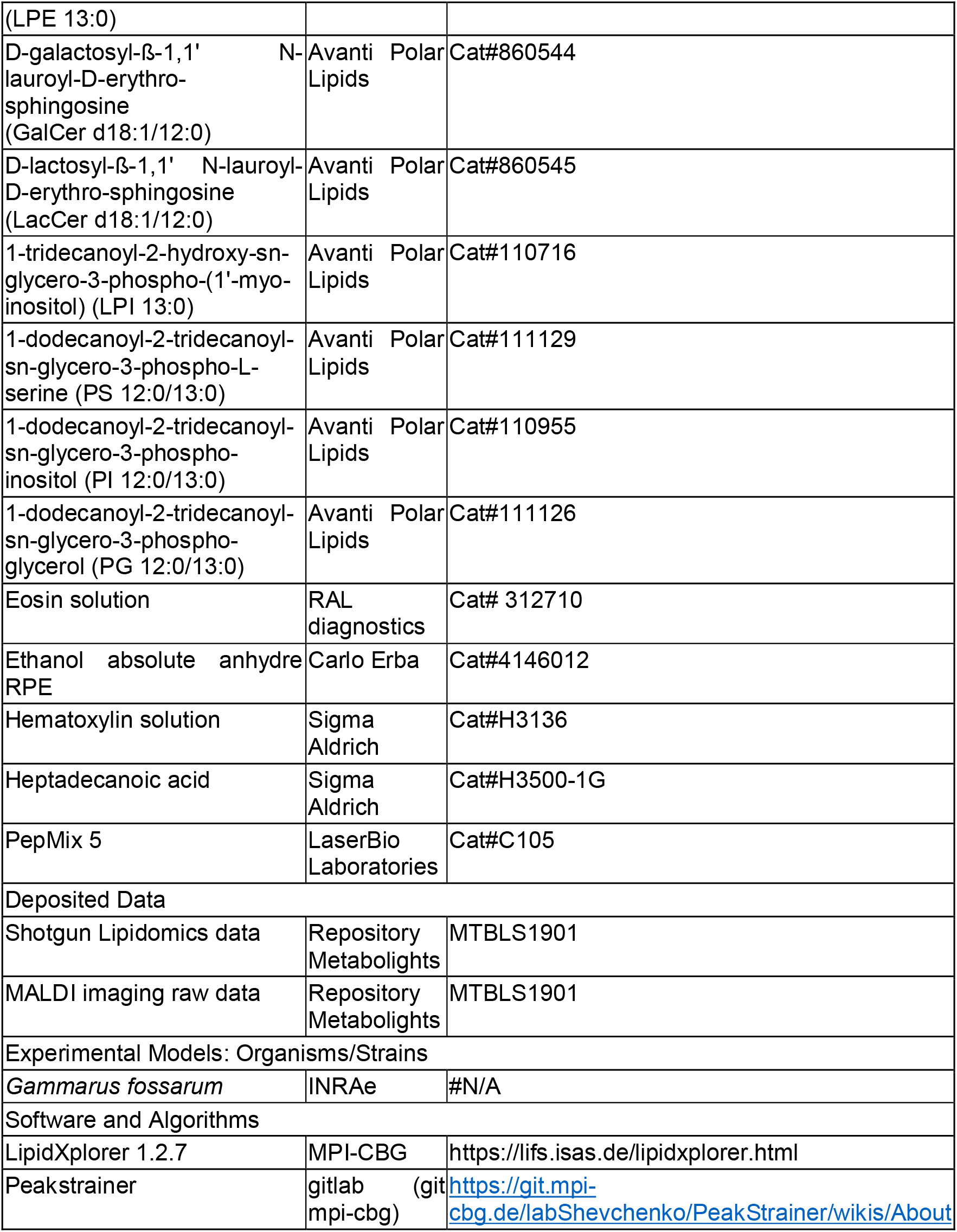
KEY RESOURCES TABLE.

### Contact for reagent and resource sharing

Further information and requests for resources and reagents should be directed to the Lead contact, Sophie Ayciriex

### Experimental model and subject details

*Gammarus fossarum* organisms were collected in a watercress site in the vicinity of the Pollon river (45°57’25.8’’N 5°15’43.6’’E) in France from a source population commonly used in our laboratory. Organisms were collected by kick sampling using a net and selected in the field according to their size by using a series of sieves (±1 cm in length). Organisms were quickly transported to the laboratory and kept in 30 L tanks supplied with continuous drilled groundwater under constant aeration without food supply. The temperature was kept at 12±1°C. After 24h, adult gammarids were sorted out at specific reproductive stages according to morphological criteria (Geffard, et al., 2010). Male gammarids in amplexus were collected and females were collected at C1 stage and D1 stage, respectively. The sampling was performed at the same time of day. All the collected gammarids were washed with deionized water, weighted and flash frozen in liquid nitrogen.

### Method details

#### Lipid extraction procedure

Entire adult gammarid (n=10 per stage) was homogenized in 300 μL of cold isopropanol with one stainless steel bead (Ø 4 mm). Protein determination assay from the homogenates was performed with BCA protein assay. ~50μg of total protein content was extracted according to a modified version of the MTBE lipid extraction procedure (Matyash et al., 2008). Briefly, 700 μL of solvent mixture (MTBE/MeOH, 10:3, *v/v*) containing one synthetic internal standard representative for each lipid class was added to the dried homogenates. Samples were vortexed for 1 h at 4°C. Phase separation was produced by adding 140 μL of water and agitating for 15 min at 4°C, followed by centrifugation (15 min, 13 400 rpm at 4°C). The upper organic phase was collected, dried down and reconstituted in 600 μL of solvent mixture CHCl_3_/MeOH, 1:2 (*v/v*). 10 μL of total lipid extract was diluted with 90 μL of solvent mixture IPA/MeOH/CHCl_3_, 4:2:1 (*v/v/v*) containing 7.5 mM ammonium formate for high resolution MS analysis.

#### Shotgun lipidomics experiment

Shotgun lipidomics analyses were performed on a Q Exactive instrument (Thermo Fischer Scientific, Bremen, Germany) equipped with a robotic nanoflow ion source TriVersa NanoMate (Advion BioSciences, Ithaca, NY, USA) using nanoelectrospray chips with a diameter of 4.1 μm. The ion source was controlled by the Chipsoft 8.3.1 software (Advion BioSciences). Ionization voltage was + 0.96 kV in positive and − 0.96 kV in negative mode. Backpressure was set at 1.25 psi in both modes by polarity switching according to Schuhmann, et al. (Schuhmann, et al., 2012). The temperature of the ion transfer capillary was set to 200°C and S-lens RF level 50%. FTMS spectra were acquired within the range of *m/z* 400–1000 from 0 min to 1.5 min in positive and then within the range of *m/z* 350–1000 from 4.2 min to 5.7 min in negative mode at a mass resolution of 140,000 (at *m/z* 200), automated gain control (AGC) of 3 × 10^6^ and with a maximal injection time of 3000 ms. Free cholesterol was quantified by parallel reaction monitoring FT MS/MS within runtime 1.51 to 4.0 min in positive ion mode. For FT MS/MS (PRM method), the micro scans were set to 1, isolation window to 1 Da, stepped normalized collision energy to 15, 25, 35, AGC to 2 × 10^4^ and maximum injection time to 650 ms. All acquired data was filtered by PeakStrainer that can be found in gitlab https://git.mpi-cbg.de/labShevchenko/PeakStrainer/wikis/About (Schuhmann, et al., 2017). Lipids were identified by LipidXplorer 1.2.7 software (Herzog, et al., 2012). Molecular Fragmentation Query Language (MFQL) queries were compiled for TAG, DAG, Cholesterol, CE, SM, PC, PC O-, LPC, LPC O-, PE, PE O-, LPE, PI, LPI lipid classes. The identification of the lipid class relied on accurately determined intact lipid masses (mass accuracy better than five ppm). The identification of the lipid molecular species relied on the MS/MS spectra inspection in both polarities of the polar head group fragments and fatty acid moieties. Lipids were quantified by comparing the isotopically corrected abundances of their molecular ions with the abundances of internal standards of the same lipid class. The amount of lipids per animal was calculated and normalized to the fresh mass of gammarids.

#### Fatty acid profiling by GC-FID

After lipid extraction of adult gammarids (n=24), lipids were dried down and reconstituted in 1 mL of methanol with 2.5% sulfuric acid containing 5 μg of heptadecanoic acid methyl ester as internal standard. The mixture was incubated at 80°C for 1 h. Then, 1.5 mL of water was added. Fatty acid methyl esters (FAMES) were extracted using 750 μL of hexane and separated in a 15 m × 0.53 mm Carbowax column (Alltech Associates, Deerfield, IL, U.S.A.) on a GC-FID (Hewlett–Packard 5890 series II). The oven temperature was programmed for 1 min at 160°C, followed by a 20°C per min ramp to 190°C and a second ramp of 5°C per min to 210°C, and then maintained at 210°C for a further 6 min. FAMES retention times were determined by comparison with those of standards and quantified using heptadecanoic acid methyl ester as standard.

#### Annotation of lipid molecular species

Annotation of lipid species follows the guidelines established by the Lipid Maps Consortium (Fahy, et al., 2005; Fahy, et al., 2009) and the Lipidomics Standards Initiative (LSI) (Liebisch, et al., 2013; Pauling, et al., 2017). Lipid species were noted as the following: <lipid class><number of total C number in the sum of fatty acid moieties>:<number of unsaturation in the sum of fatty acid moieties> (e.g., PC 34:1). Lipids molecular species were annotated as the following: <lipid class><number of C in the first fatty acid moiety>:<number of double-bond in the first fatty acid moiety>–<number of total C number in the second fatty acid moiety>:<number of double bond in the second fatty acid moiety> [e.g., phosphatidylcholine (PC) 16:0–18:1]. Sphingolipid species were annotated as <lipid class><number of C in the long-chain base (LCB) + fatty acid moieties>:<number of double bonds in the LCB + fatty acid moieties> (e.g., SM 34:1).

#### Preparation of gammarid tissue sections

The male and female gammarids of C1 and D1 stages were directly plunge-frozen in liquid N_2_ without embedding and then stored at −80°C until cryo-sectioning. Transversal and longitudinal sections of adult *G. fossarum* were cut at −23°C with a thickness of 12 μm utilizing a MICROM HM505E cryostat microtome. The sections were immediately thaw mounted onto ITO coated glass slides (Sigma-Aldrich) and dried for 30 min in a desiccator under low vacuum. Then the slides were placed in plastic bags filled with N_2_ to avoid oxidation and stored at −80°C until analysis.

All optical images were recorded at 10X magnification on an Olympus BX41M optical microscope.

#### MALDI-TOF imaging

The longitudinal sections of male gammarid were coated with a homogeneous layer of matrices using a robotic TM-Sprayer (HTX Technologies, Chapel Hill, NC, USA) prior to MALDI imaging. 10 mg/mL CHCA solution prepared in ACN/H_2_O/TFA (70:30:0.1, *v/v/v*) and 10 mg/mL 9-AA solution prepared in EtOH/H_2_O (70:30, *v*/*v*) were used for positive and negative ion mode MALDI imaging experiments, respectively. CHCA solution was sprayed at 70°C and 9-AA solution at 90°C with the following parameters: flow rate: 0.12 mL/min; nozzle height: 40 mm; nozzle moving speed: 120 cm/min; moving pattern: CC; track spacing: 3 mm; drying time: 30 s; nebulizing gas (N_2_) pressure: 10 psi; 2 passes. MALDI imaging experiments were carried out with an UltrafleXtreme MALDI-TOF/TOF mass spectrometer (Bruker Daltonics, Wissembourg, France) equipped with a 2 kHz Smart beam-II™ Nd:YAG laser (wavelength: λ=355 nm). The ion images were acquired with a pixel size of 40 μm (‘medium’ focus setting) and the spectrum of each pixel represented ion signals summed from 500 laser shots. The mass spectra were acquired in reflectron mode over a mass range of *m/z* 120 to 1700. Mass calibration was achieved using calibration standard PepMix 5 (LaserBio Labs, Sophia Antipolis, France) which was deposited onto the matrix coated slide. Data acquisition and processing were performed using flexControl 3.4 and flexImaging 4.1 (Bruker Daltonics, Wissembourg, France), respectively. ‘Median’ method was employed for normalization of all the mass spectra given that it provided slightly higher signal to noise ratio.

#### TOF-SIMS imaging

TOF-SIMS imaging experiments were performed on a TOF-SIMS V (IONTOF GmbH, Münster, Germany) mass spectrometer. The bismuth liquid metal ion gun (LMIG) was operated in high current bunch (HCBU) mode to ensure a good mass resolution and a sufficient beam current together with a reasonable analysis time. The 25 keV Bi_3_^+^ cluster ions were selected as primary ion beam, of which the current was about 0.45 pA measured at 10 kHz with a fixed pulse width of 20.5 ns. The secondary ions were extracted and accelerated to 2 keV at the entrance of the TOF analyzer, and then post-accelerated to 10 keV before reaching the detector which is composed of a single micro-channel plate, a scintillator and a photomultiplier. A low energy pulsed electron flood gun (20 eV) was employed to compensate the charge accumulation on the insulating tissue samples. Ion images were generated from areas of 500 μm × 500 μm divided by 256 × 256 pixels. The total ion dose applied on each area is ~ 5×10^11^ ions/cm^2^. Data processing was performed using SurfaceLab 7 software (IONTOF GmbH, Münster, Germany). Mass spectra were internally calibrated using small fragments commonly observed in SIMS spectra such as CH^+^, CH_2_^+^, CH_3_^+^, C_2_H_3_^+^, C_2_H_5_^+^ in positive ion mode and CH^−^, CH_2_^−^, C_3_^−^, C_4_^−^, C_4_H^−^ in negative ion mode. Improvement of mass accuracy was obtained by adding characteristic ion peaks of Vitamin E (*m/z* 429.373, C_29_H_49_O_2_ ^−^ and *m/z* 430.381, C_29_H_50_O_2_·^+^) to the mass calibration list.

#### Histological staining

The post MSI analyzed tissue sections were stained with Hematoxylin and Eosine (H&E) to visualize the anatomy of the tissue sections. After washing away the MALDI matrices with ethanol, the tissue sections were stained in hematoxylin solution for 15 min before being washed with running tap water for 5 min. Then the sections were immerged in eosin solution for 10 min. After washing the tissues with distilled water for 5 min, the slides were placed successively in 70% ethanol, 96% ethanol and 100% ethanol, each for 2 min. Finally, the slides were plunged in n-butanol solution for 4 min and then allowed to dry at ambient atmosphere before observation under a microscope.

#### MS/MS analysis of unknown sulfate-based lipids in hepatopancreas

Hepatopancreas were dissected and pooled from 10 male gammarids for lipid extraction and subsequent MS analysis. For lipid extraction, hepatopancreas tissues were homogenized in 300 μL methanol with zirconia beads (Ø 0.5 mm) and then extracted with a mixture of MeOH/CHCl_3_ (1/2, *v*:*v*) through continuous agitation at 4°C for 1 h. 100 μL H_2_O were added for phase separation. After a quick centrifugation, the lower phase containing the lipids was collected and dried down at 40°C under constant N_2_ flow. The final product was reconstituted in pure methanol for MS analysis.

The lipid extract was infused directly into the HESI source of a Q Exactive mass spectrometer (Thermo Fischer Scientific, Bremen, Germany) with a Hamilton™ syringe (1 mL, 600 μL/min). For both MS and MS/MS analyses, the spray voltage was set at + 4 kV for positive ion mode and −3.3 kV for negative ion mode. Capillary temperature was 320°C and S-lens level was 75. For MS/MS fragmentation, the AGC target was set at 5 × 10^6^ and maximum injection time was 100 ms. Collision energy used to fragment the selected precursors was optimized and set at a normalized value of 50 except for the precursor ions at *m/z* 295.212 and *m/z* 297.1369, for which the normalized collision energy were 45 and 40, respectively. The mass resolution was 140,000 for all the analyses.

## Data and software availability

Shotgun lipidomics data and MS imaging data have been deposited to the EMBL-EBI MetaboLights database (Haug, et al., 2020) with the identifier MTBLS1901. The datasets can be accessed here: https://www.ebi.ac.uk/metabolights/MTBLS1901.

## Acknowledgments

This work was supported by the French National Research Agency (ANR) (young investigator grant, ANR-18-CE34-0008 PLAN-TOX), the Chemistry Institute of Lyon (young investigator inter-laboratory cooperation grant, ICL-2017 DOSAGE), the ISA research funds grant, and also the French GDR “Aquatic Ecotoxicology” framework which aims at fostering stimulating discussions and collaborations for more integrative approaches. MALDI-TOF instrument at CNRS-ICSN was founded by the Région Ile-de-France (DIM Analytics). We would like to thank Elodie Chauvet, Yves de Puydt from Tescan Analytics (Fuveau, France) for the access to TOF-SIMS instrument and the help with the experiments. We thank Jean-Valery Guillaubez for his technical assistance on the lipid MS analysis of the hepatopancreas sample. We would also like to warmly thank Dr. Serge Della-Negra for his support and fruitful discussions. We thank Dr. Alain Brunelle for his careful and critical reading of the manuscript. The authors thank Nicolas Delorme, Laura Garnero, Hervé Quéau for their technical assistance in gammarids sampling and Guy Charmantier for his help on gammarids histology. We thank the Pôle Biosciences, Technologies, Ethique of Lyon Catholic University for the access to the microtome.

## Author Contributions

T.F., O.G., A.C., D.D.E., S.A. designed research; T.F., O.K., E.T., N.E., S.A. performed the experiments; J.L., A.S., A.Sh., D.T. contributed analytic tools; T.F., Y.C., S.A. performed data processing; T.F., O.K., O.G., Y.C., E.T., S.A. analyzed data; T.F., O.G. and K.A. conducted histology analysis; S.A. provided funding, project administration and resources; and T.F, S.A. prepared the figures and wrote the manuscript. All authors have provided feedback on the manuscript and have approved the final version.

## Declaration of Interests

The authors declare no competing interests.

## Additional information

Supplementary information: Figures S1 – S19 and Table S1.

## Notes

https://www.ebi.ac.uk/metabolights/MTBLS1901

### Competing Interest Statement

The authors have declared no competing interest.

