## Supplementary information for "Shotgun lipidomics and mass spectrometry imaging unveil diversity and dynamics in lipid composition in *Gammarus fossarum*"

Figure S19. Defining the oocyte areas by comparing the optical image with H&E stained images………………………………………………………………………………………………...17

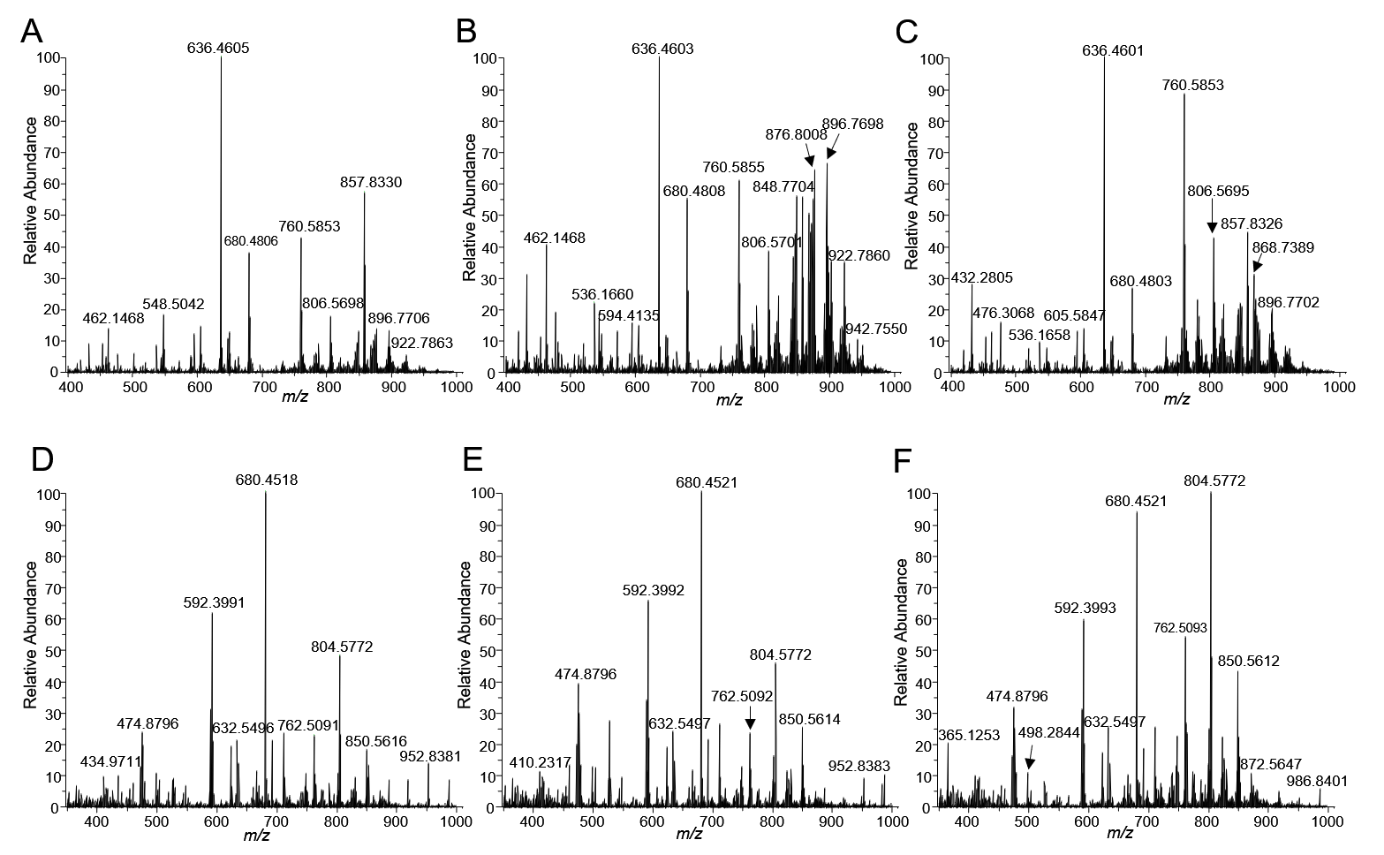

**Figure S1. Representative mass spectra of G. fossarum lipid extract**. Spectra were collected in m/z range 400-1000 in positive ion mode (A-C) and in negative ion mode (D-F). A. Mass spectra of lipid crude extract from female at C1 stage. B. Mass spectra of lipid crude extract from female at D1 stage. C. Mass spectra of lipid crude extract from male. D. Mass spectra of lipid crude extract from female at C1 stage. E. Mass spectra of lipid crude extract from female at D1 stage. F. Mass spectra of lipid crude extract from male.

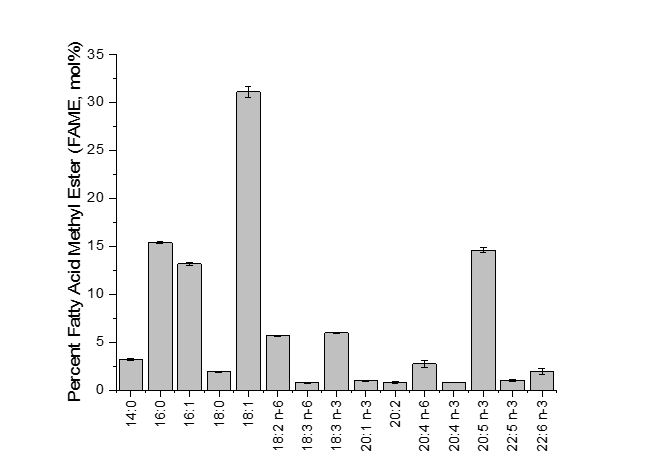

**Figure S2. Fatty acid composition in total G. fossarum lipid extract**. FA composition was determined using GC-FID from adult organisms (n=24). The prominent FA is oleic acid (18:1) followed by palmitic acid (16:0) and eicosapentaneoic acid (20:5). n-3 (omega-3) symbol represents the presence of a double bond three C atoms away from the terminal methyl group in the chemical structure.

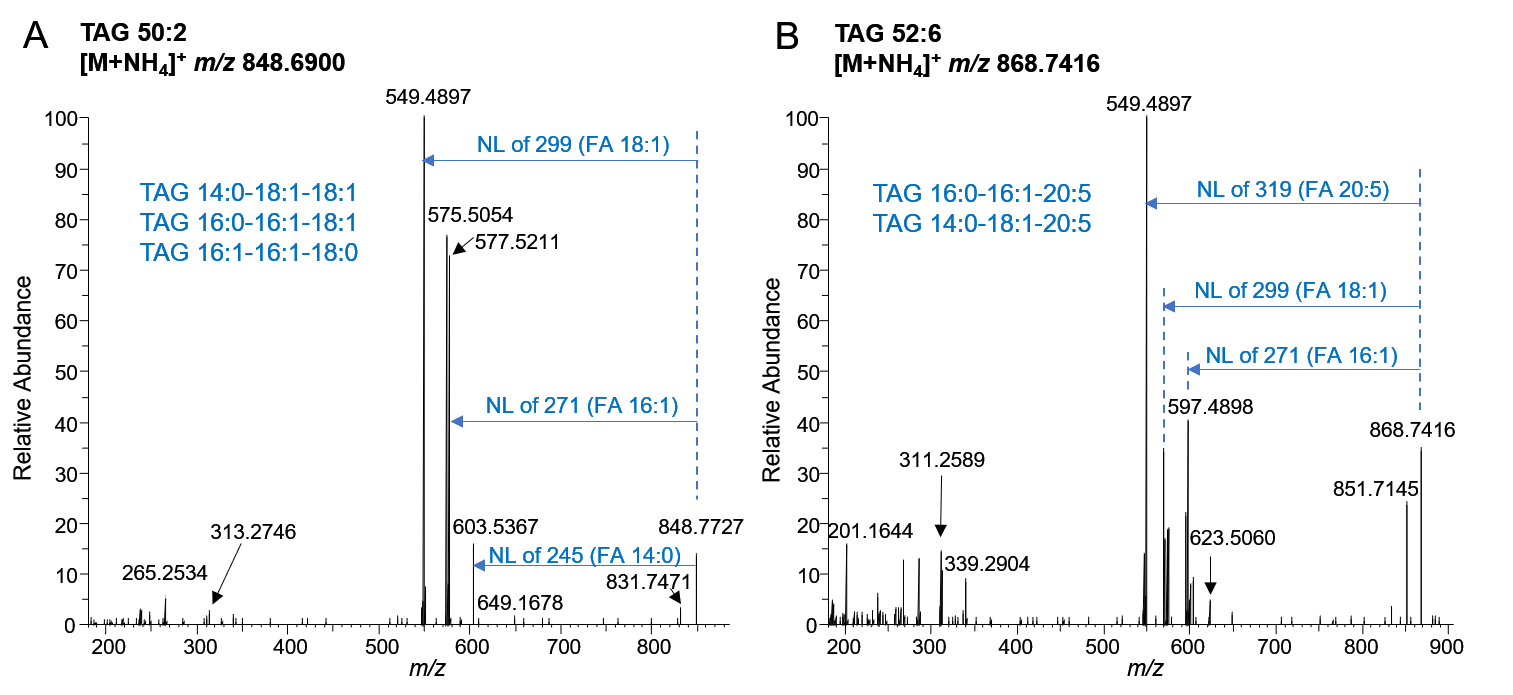

**Figure S3. MS/MS spectra of major TAG species acquired in positive ion mode (HCD=25).** (A) MS/MS spectrum of TAG 50:2 precursor ion, exhibiting the following neutral loss (NL): 299, 271 and 245. (B) MS/MS spectrum of TAG 52:6 precursor ion, exhibiting NL of 319, 271, 299 and 245.

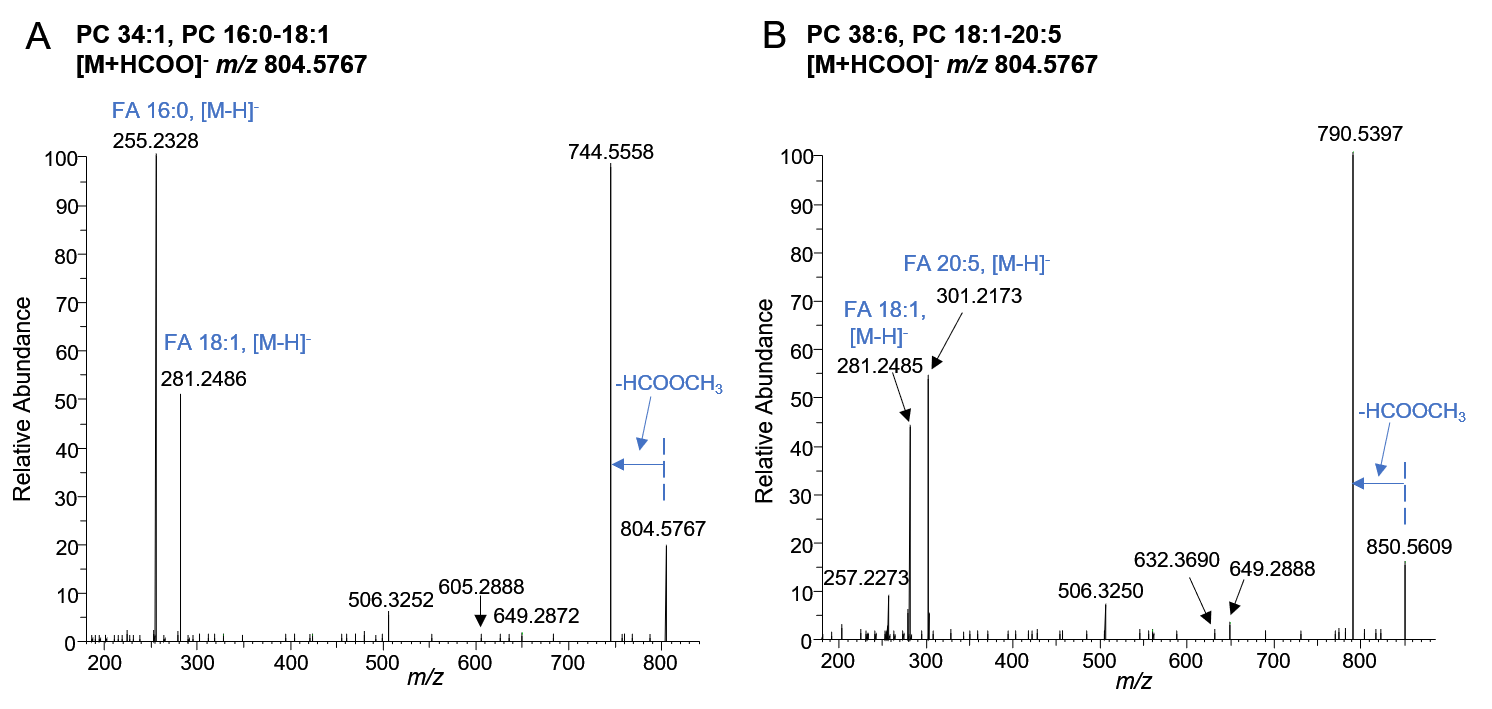

**Figure S4. MS/MS spectra of the major PC species acquired in negative ion mode (HCD=25).** PC species were detected in negative ion mode as formate adduct. (A) MS/MS spectrum of PC 34:1 precursor ion at m/z 804.5767 which was annotated as PC 16:0-18:1. (B) Mass spectrum of PC 38:6 precursor ion at m/z 850.5609 which was annotated as PC 18:1-20:5.

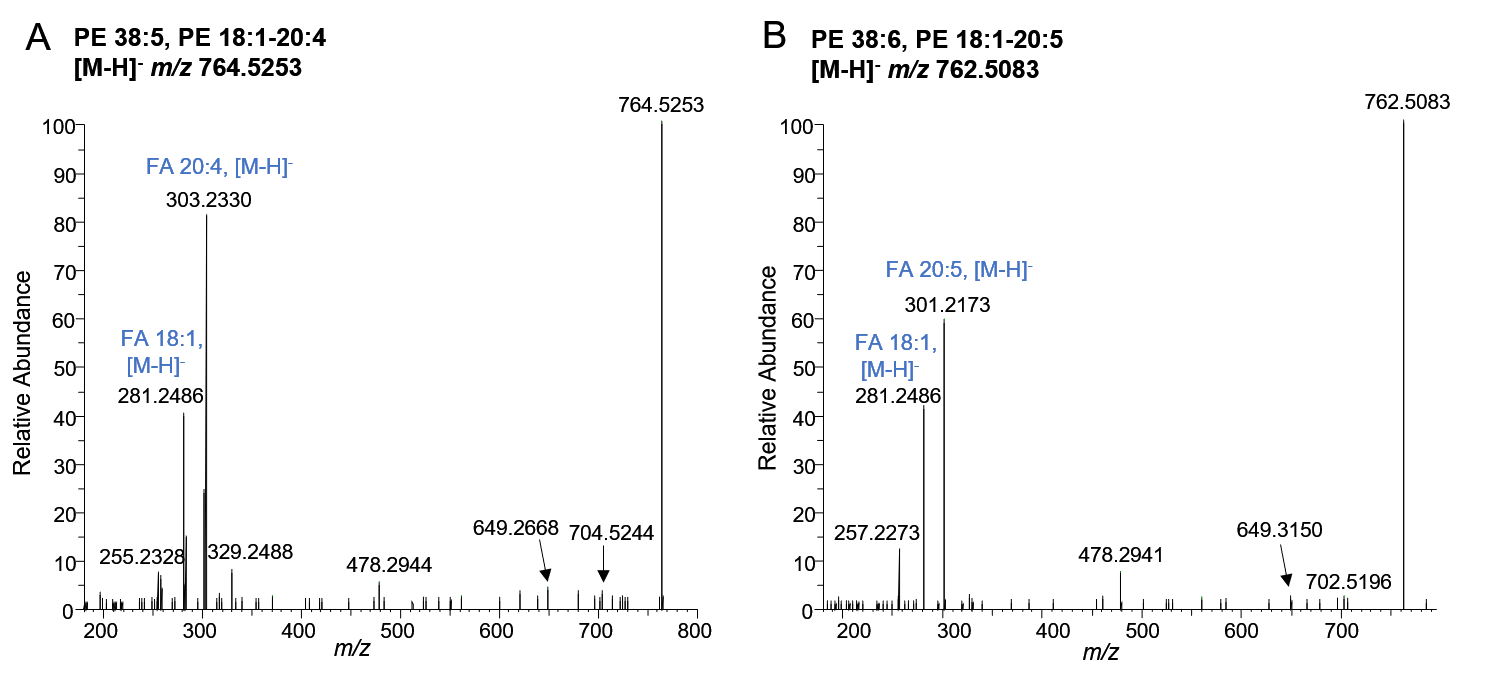

**Figure S5. MS/MS spectra of the major PE species acquired in negative ion mode (HCD=25).** PE species were detected in negative ion mode as deprotonated molecule. (A) MS/MS spectrum of PE 38:5 precursor ion at m/z 764.5253 which was annotated as PE 18:1-20:4. (B) Mass spectrum of PE 38:6 precursor ion at m/z 762.5083 which was annotated as PE 18:1-20:5.

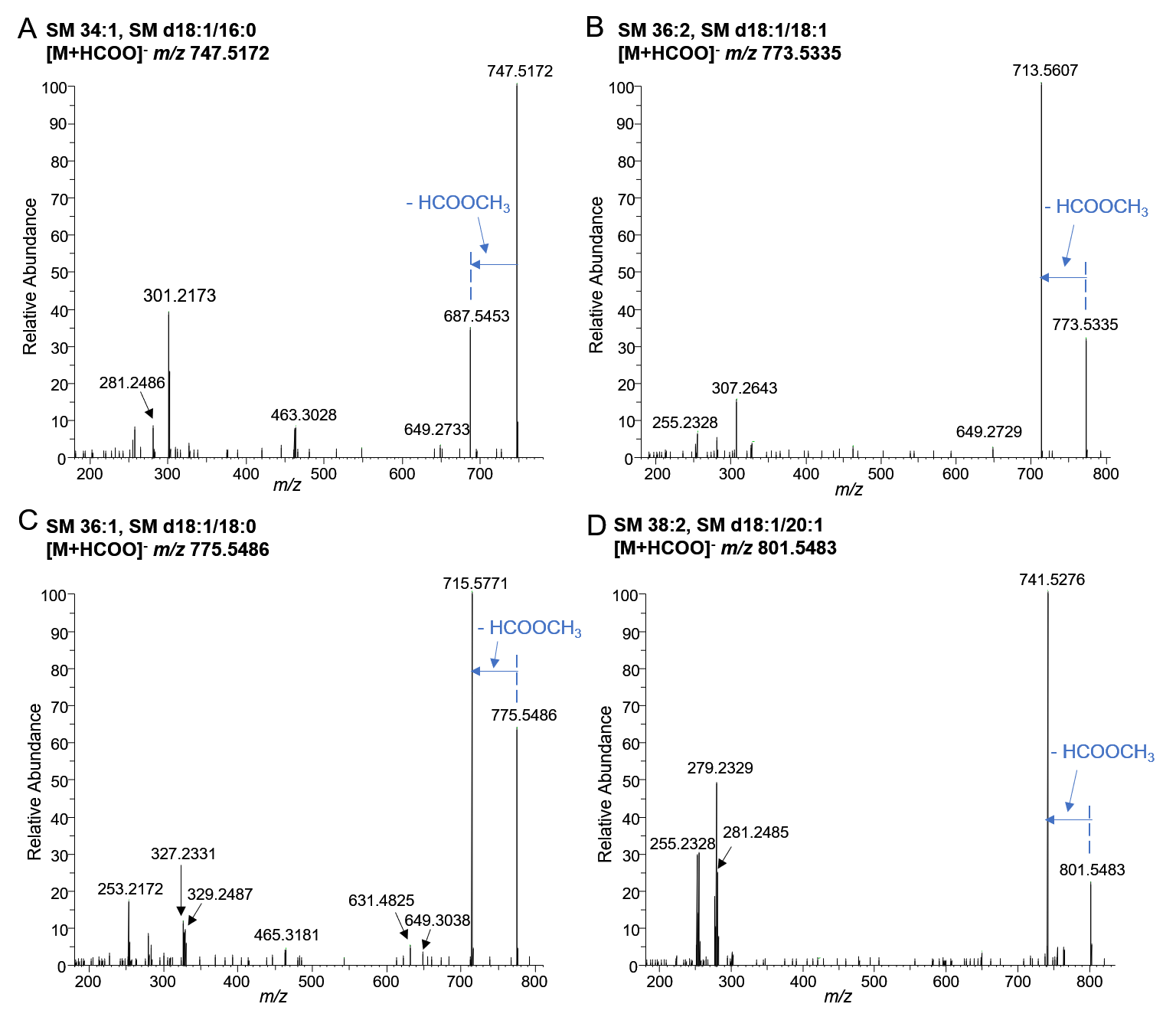

**Figure S6. MS/MS spectra of SM species acquired in negative ion mode (HCD=25). SM species were detected in negative ion mode as formate adduct.** (A) MS/MS spectrum of SM 34:1 precursor ion at m/z 747.5172. (B) MS/MS spectrum of SM 36:2 precursor ion at m/z 773.5335. (C) MS/MS spectrum of SM 36:1 precursor ion at m/z 775.5486. (D) MS/MS spectrum of SM 38:2 precursor ion at m/z 801.5483.

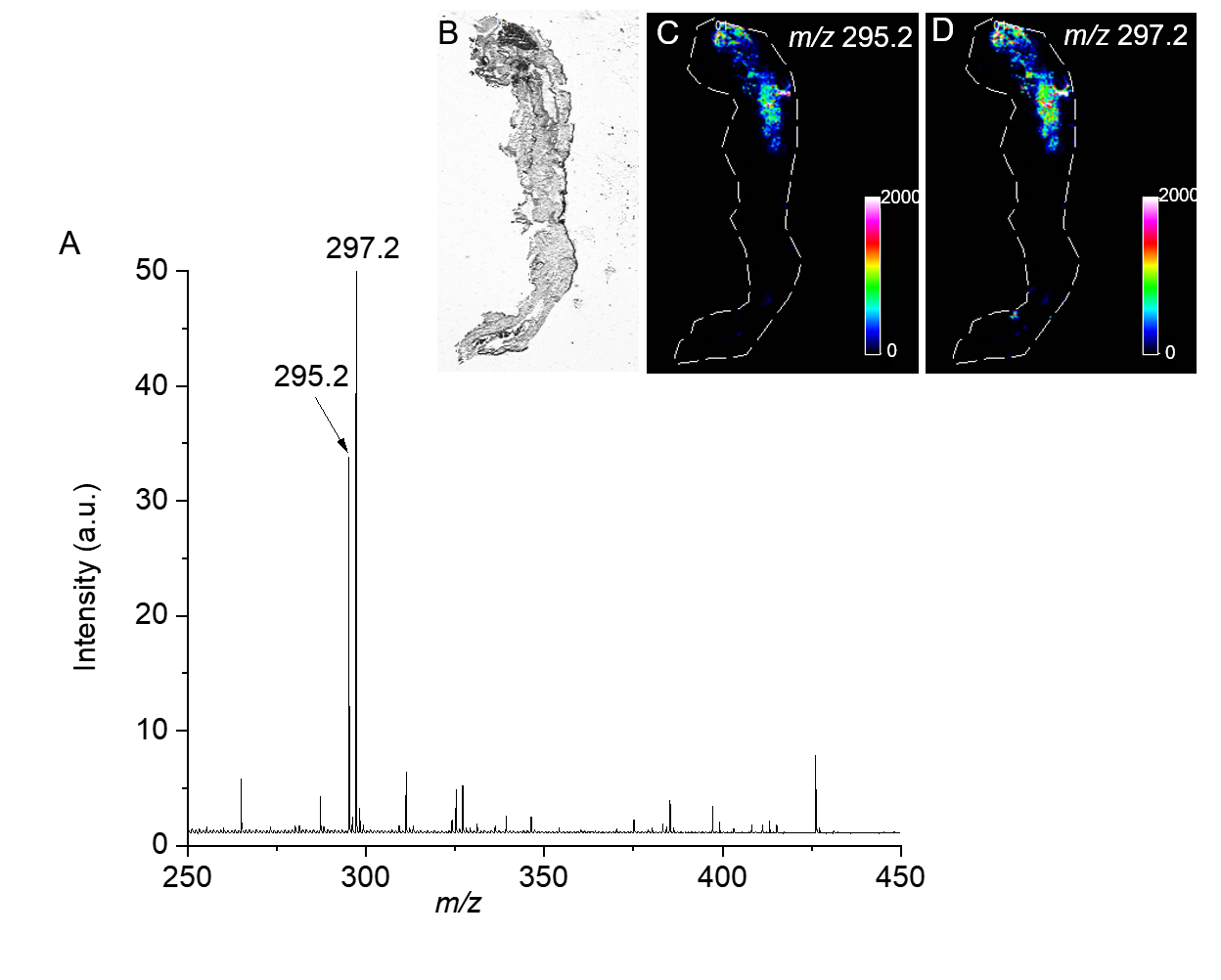

**Figure S7. MALDI MSI data acquired in negative ion mode. (A) Partial MALDI-TOF mass spectrum in the mass range of m/z 250-450.** (B) Optical image of the analyzed whole-body gammarid tissue section. (C) Ion image of the ion at m/z 295.2. (D) Ion image of the ion at m/z 297.2.

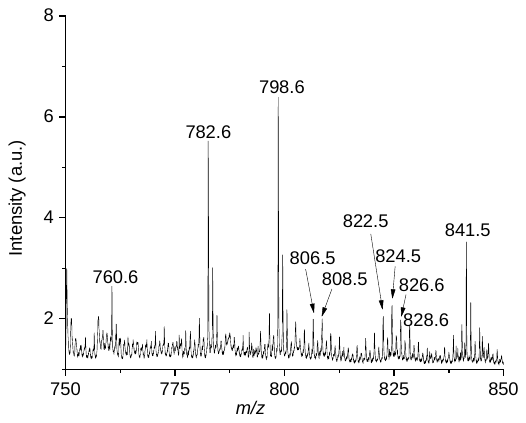

**Figure S8. Partial MALDI-TOF mass spectrum in the mass range of m/z 750-850 acquired in positive ion mode.**

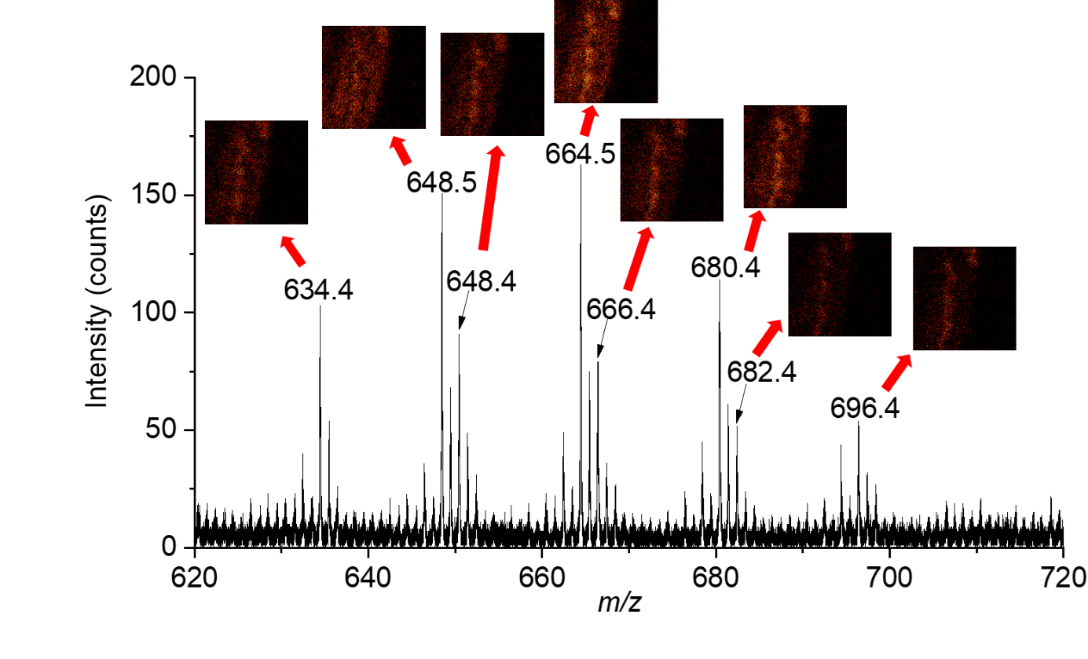

**Figure S9. Partial TOF-SIMS spectrum showing the detection of unknown ion species at m/z 634.4-696.4 in positive ion mode.** Ion images of individual ion species illustrating their localization in the hepatopancreas are also shown.

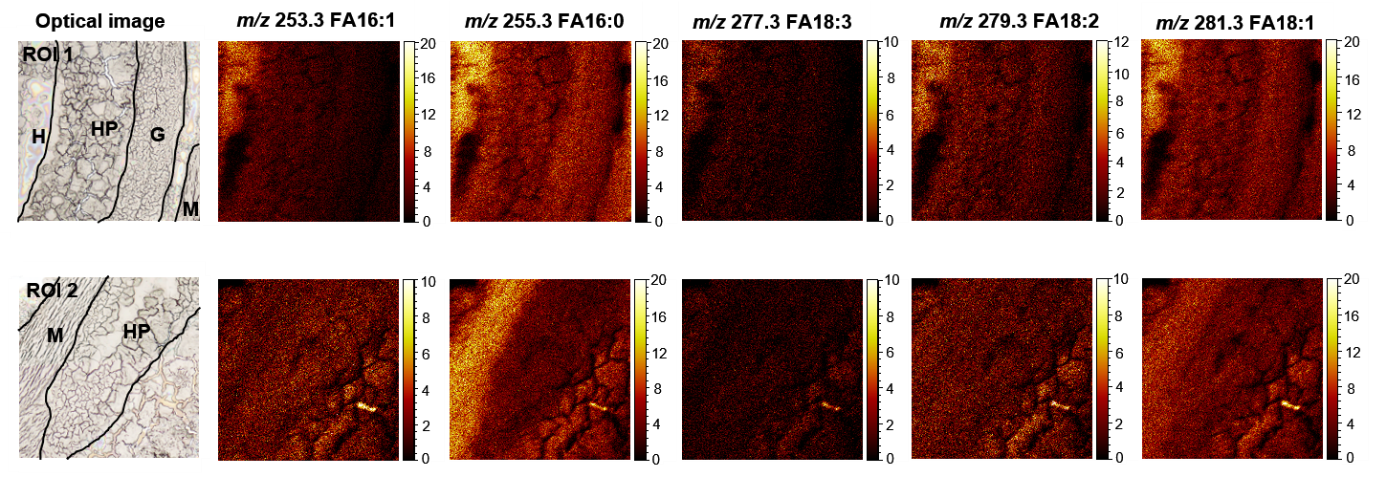

**Figure S10. Ion images of fatty acids detected from the two regions of interest.**

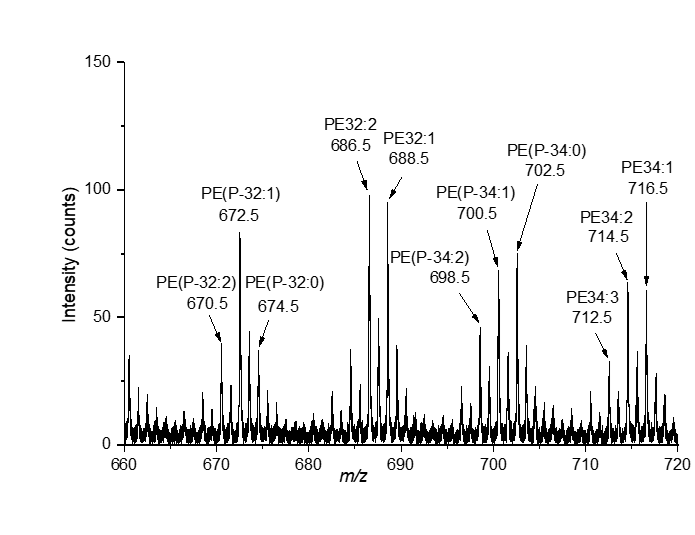

**Figure S11. Partial TOF-SIMS spectrum showing the detection of phosphatidylethanolamine (PE) lipids in the mass range of m/z 660-720 in negative ion mode.**

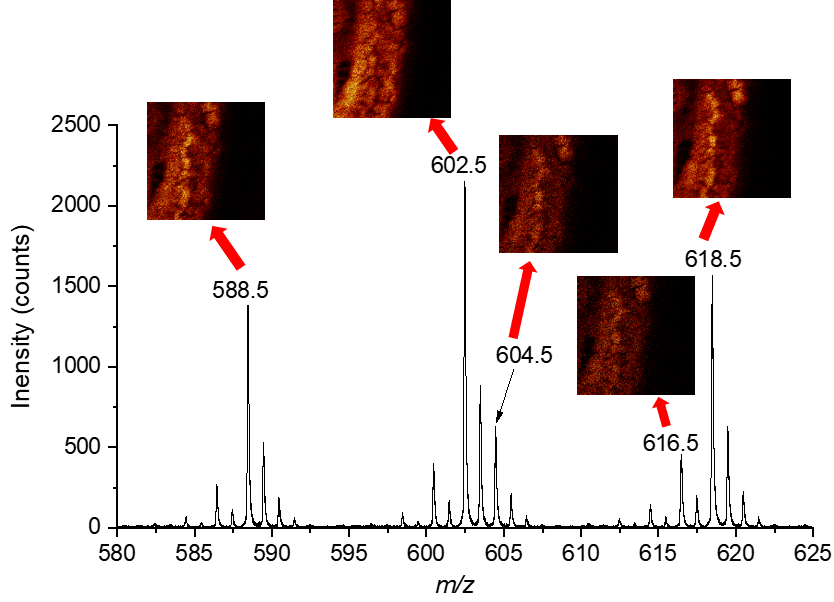

**Figure S12. Partial TOF-SIMS spectrum showing the detection of ions at m/z 588-618 in negative ion mode.** Ion images of individual ion species illustrating their localization in the hepatopancreas are also shown.

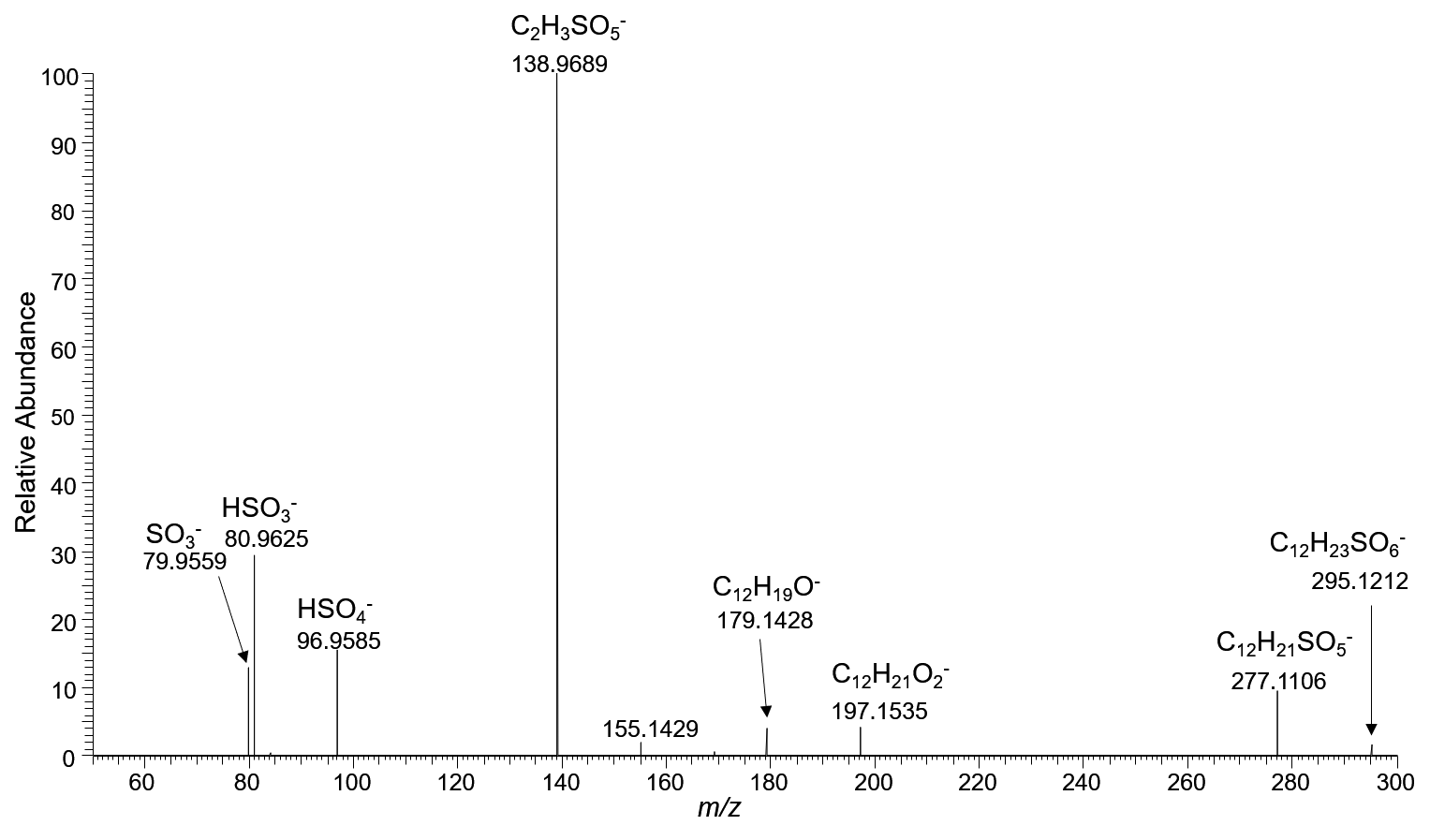

**Figure S13. Orbitrap MS/MS spectrum of the precursor ion at m/z 295.1212 from lipid extract of hepatopancreas of male gammarid.**

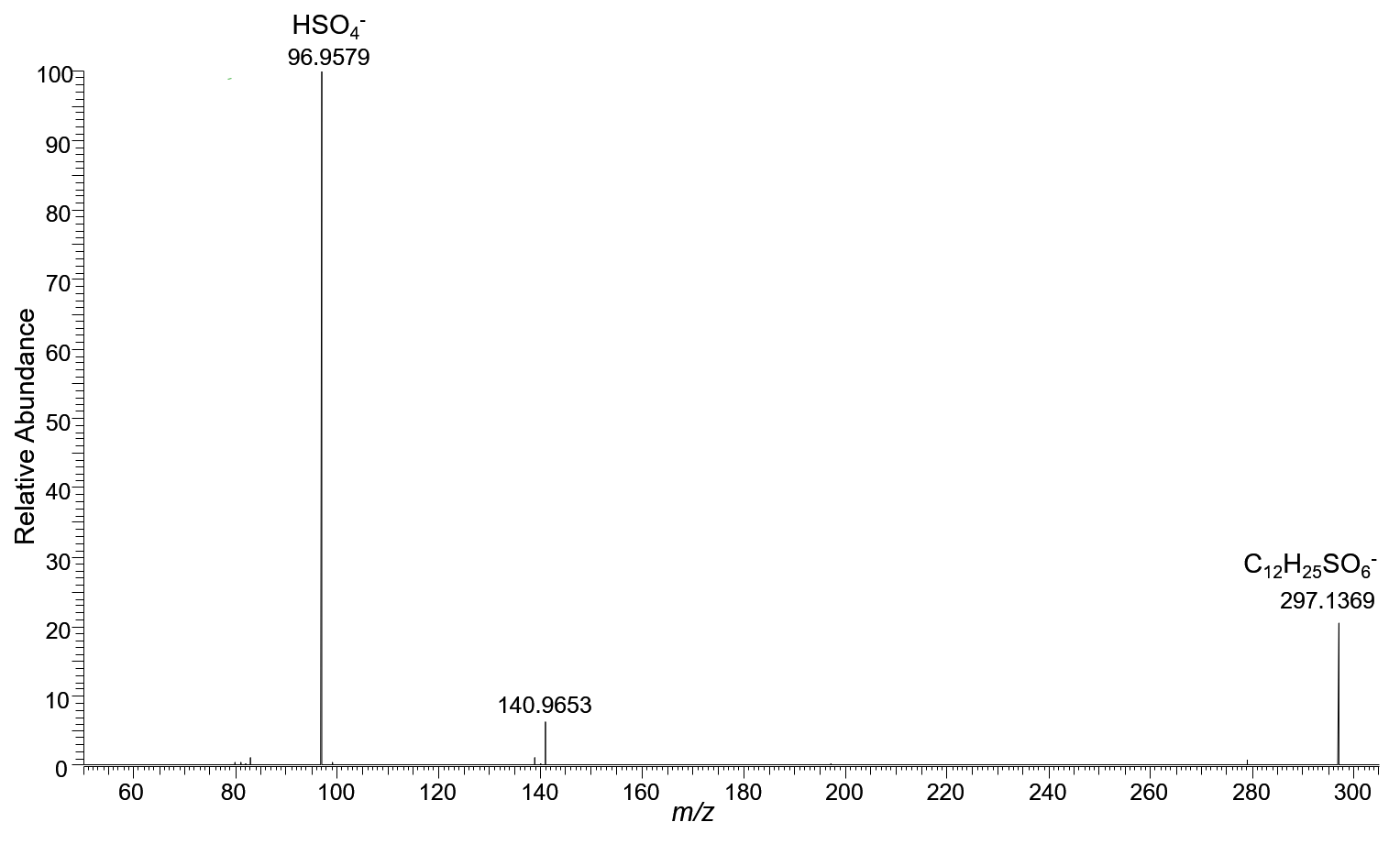

**Figure S14. Orbitrap MS/MS spectrum of the precursor ion at m/z 297.1369 from lipid extract of hepatopancreas of male gammarid.**

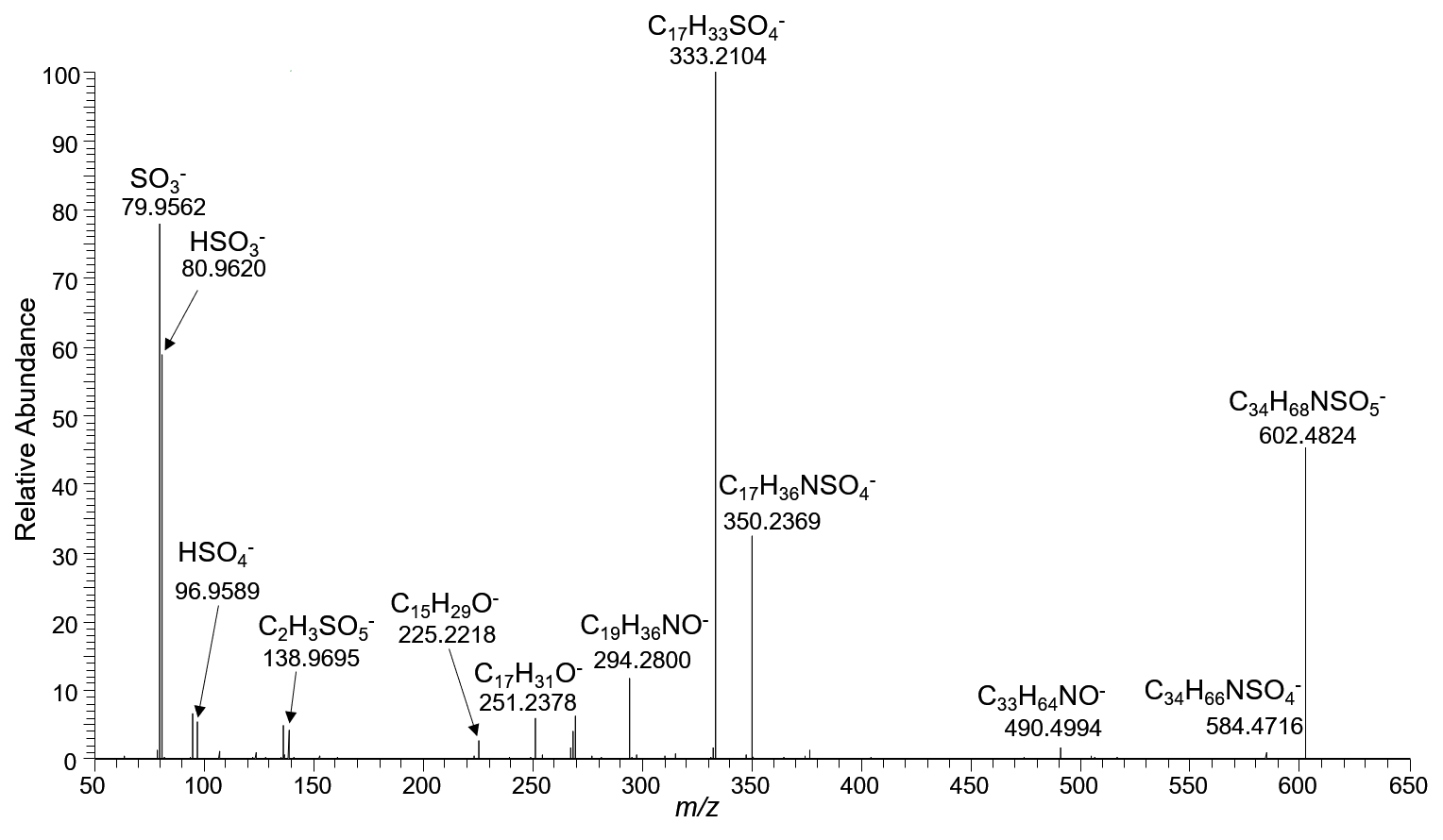

**Figure S15. Orbitrap MS/MS spectrum of the precursor ion at m/z 602.4824 from lipid extract of hepatopancreas of male gammarid.**

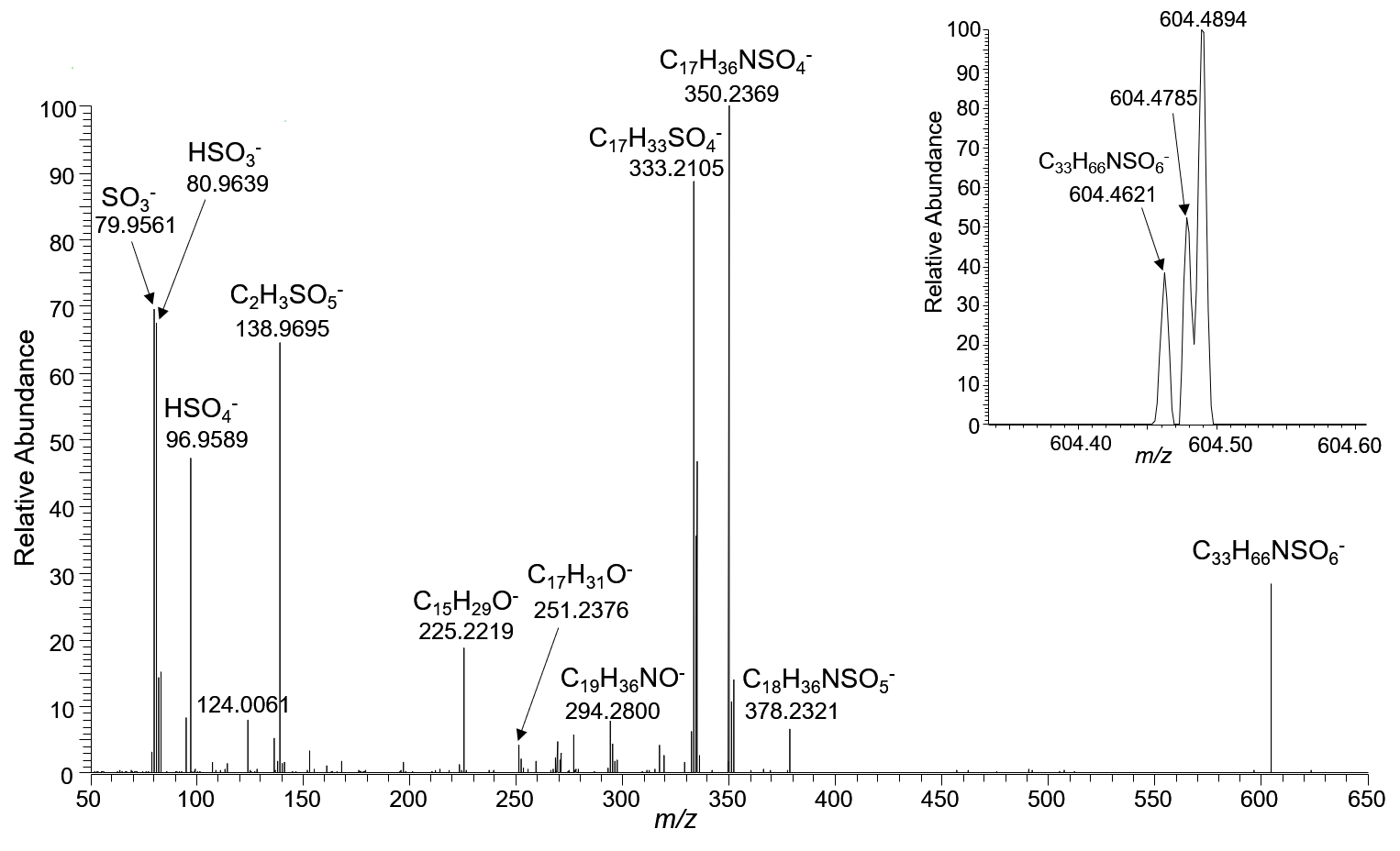

**Figure S16. Orbitrap MS/MS spectrum of the precursor ion at m/z 604.4621 from lipid extract of hepatopancreas of male gammarid.**

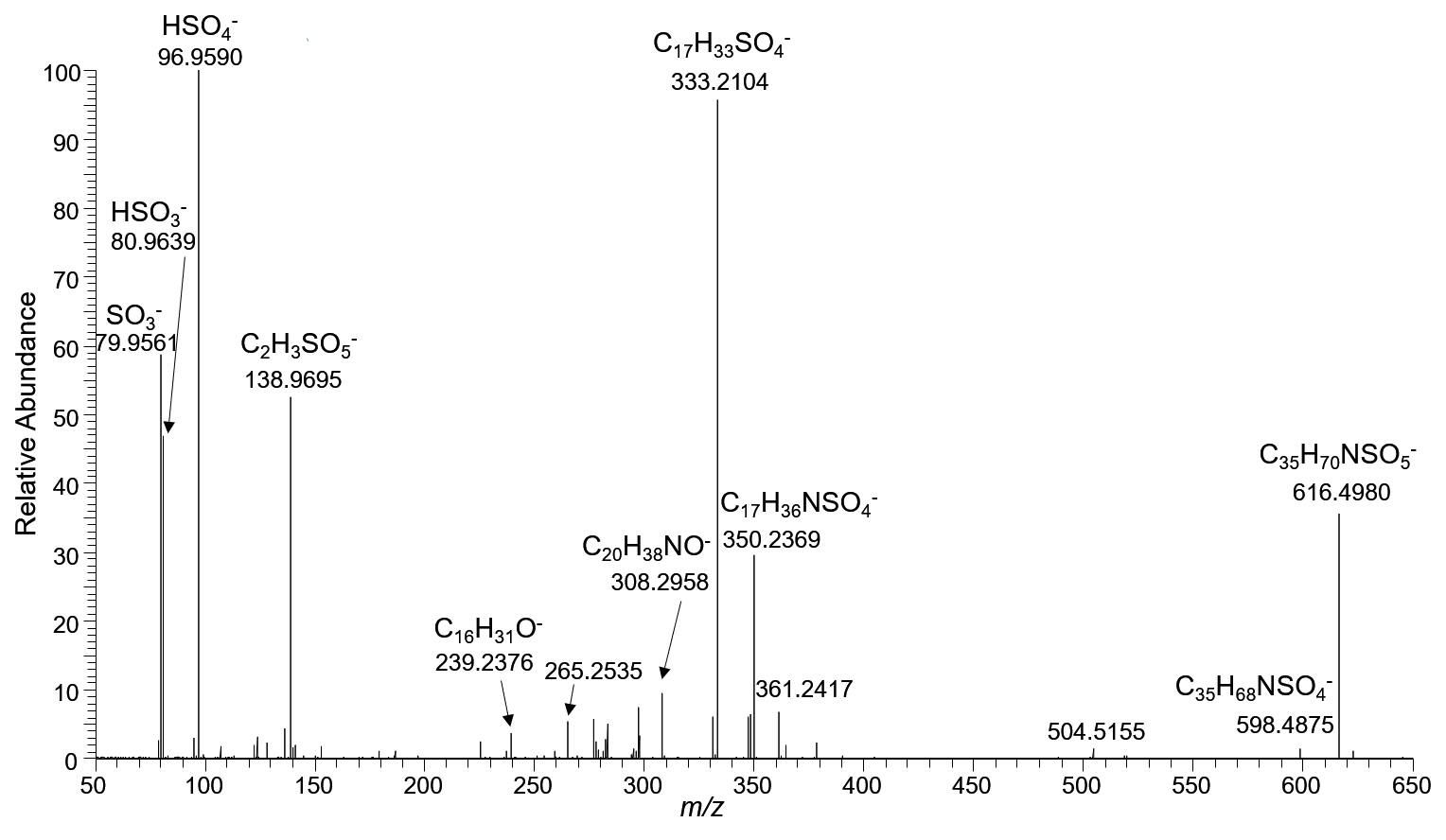

**Figure S17. Orbitrap MS/MS spectrum of the precursor ion at m/z 616.4980 from lipid extract of hepatopancreas of male gammarid.**

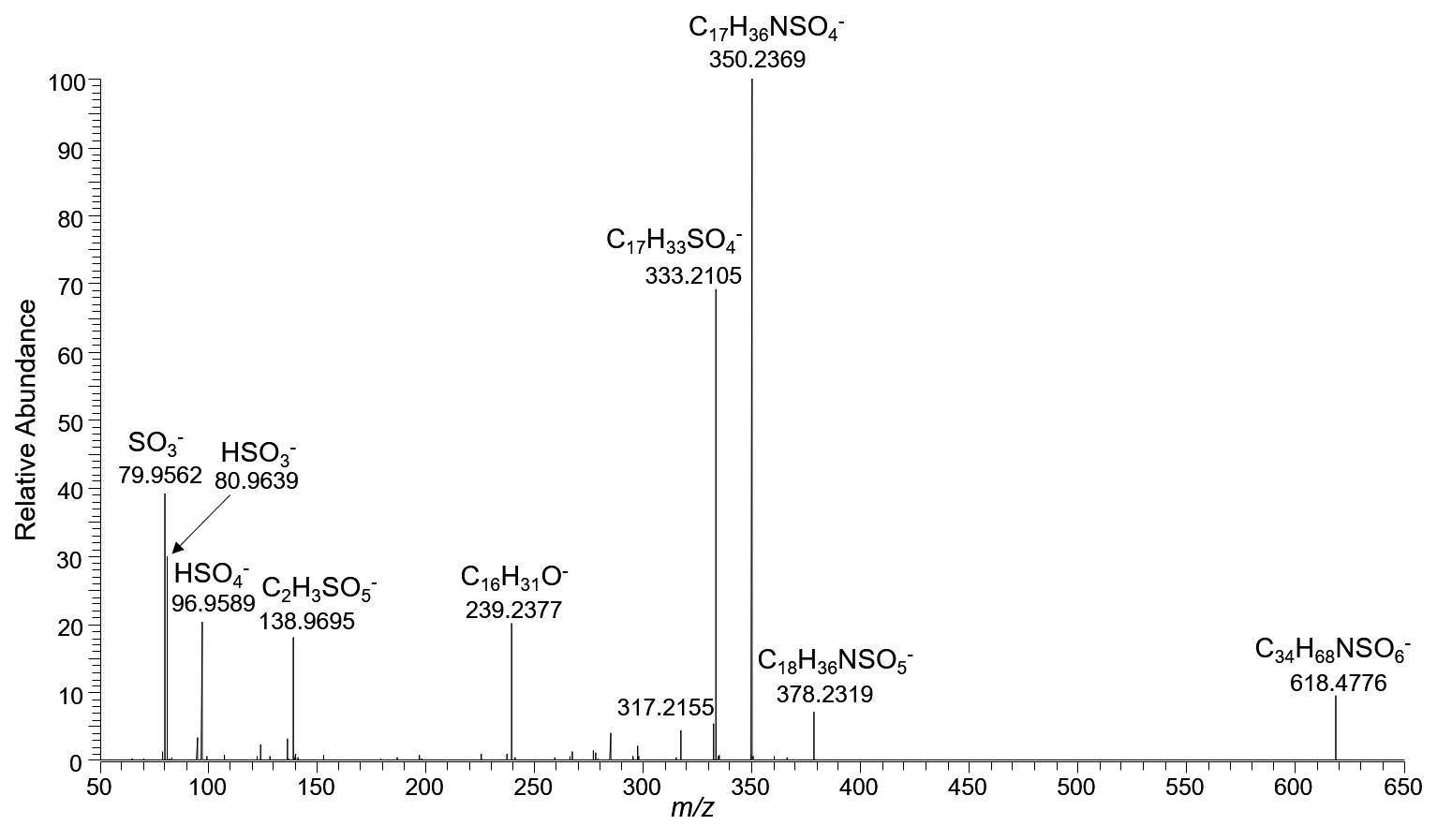

**Figure S18. Orbitrap MS/MS spectrum of the precursor ion at m/z 618.4776 from lipid extract of hepatopancreas of male gammarid.**

**Table S1. Assignments of the unknown ion species observed in MSI.**

| *m/z* (MALDI &SIMS)) | *m/z* (Orbitrap) | Assignment | Theoretical *m/z* | Mass deviation (ppm) |
| --- | --- | --- | --- | --- |
| 295.2 | 295.1212 | C_12_H_23_SO_6_^-^ | 295.1215 | -1.0 |
| 297.2 | 297.1369 | C_12_H_25_SO_6_^-^ | 297.1372 | -1.0 |
| 588.5 | 588.4667 | C_33_H_66_NSO_5_^-^ | 588.4662 | 0.85 |
| 602.5 | 602.4824 | C_34_H_68_NSO_5_^-^ | 602.4818 | 1.0 |
| 604.5 | 604.4621 | C_33_H_66_NSO_6_^-^ | 604.4611 | 1.6 |
|  | 604.4785 | Not assigned | N/A | N/A |
|  | 604.4894 | Not assigned | N/A | N/A |
| 616.5 | 616.4980 | C_35_H_70_NSO_5_^-^ | 616.4975 | 0.81 |
| 618.5 | 618.4776 | C_34_H_68_NSO_6_^-^ | 618.4767 | 1.5 |

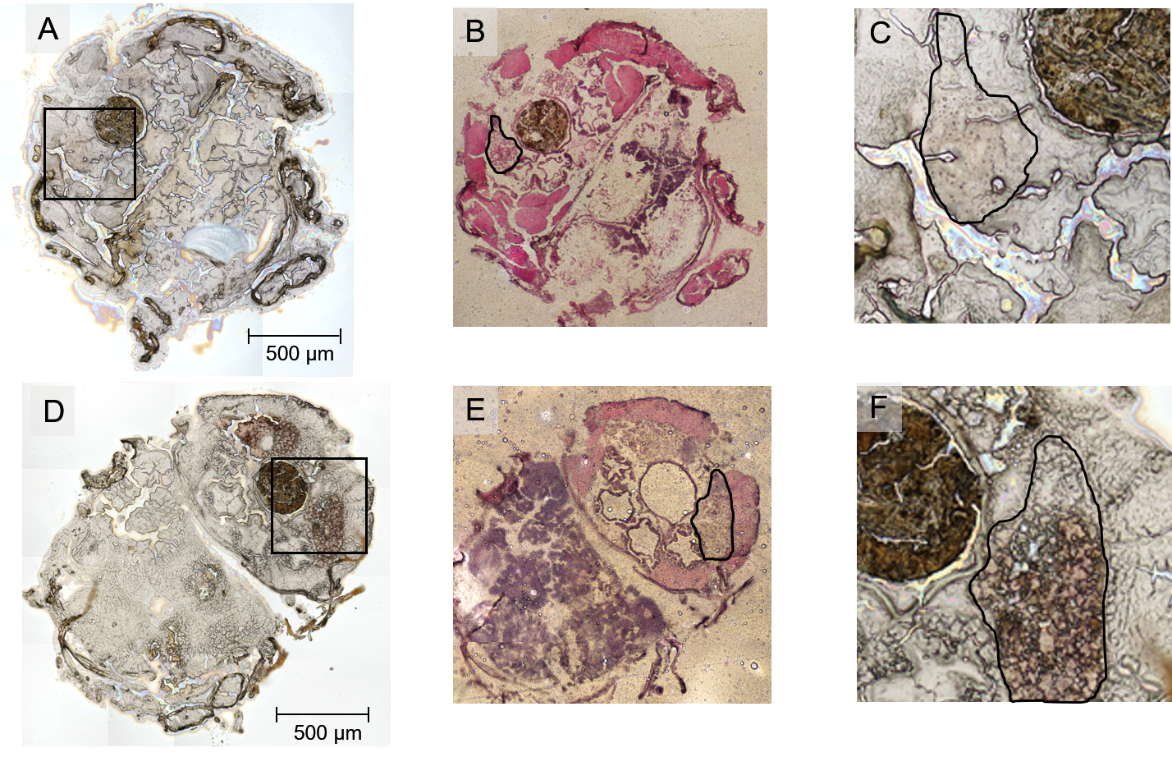

**Figure S19. Defining the oocyte areas by comparing the optical image with H&E stained images.** (A) Optical image of transvers tissue section of female gammarid at C1 stage. (B) H&E stained image of the post-analyzed tissue section of female gammarid at C1 stage. (C) Zoomed image of the analyzed area shown in (A). (D) Optical image of transvers tissue section of female gammarid at D1 stage. (E) H&E stained image of the post-analyzed tissue section of female gammarid at D1 stage. (F) Zoomed image of the analyzed area shown in (D). The black square indicates the region of interest analyzed by TOF-SIMS. The oocytes areas are outlined with black line.
